# Protein hunger-driven nitrogen flow from phospholipids to amino acids in *Drosophila*

**DOI:** 10.64898/2026.09.18.752762

**Authors:** Tianji Ma, Guangyan Wu, Lauren Tham, Madison Palarca-Wong, Sharon Vaz, Nick Allen, Carrie Shao, Ashley Tsai, Richard W. Song, Pamela M. England, Qili Liu

## Abstract

Protein is unique among macronutrients due to its nitrogen content and that its building blocks, essential amino acids, cannot be synthesized by animals. Additionally, animals lack specialized amino acid storage sites, necessitating a continuous dietary intake. In the face of insufficient dietary protein, how do animals manage nitrogen limitation? Using *Drosophila*, we reveal a previously undocumented nitrogen flow from phospholipids to amino acids, a process enhanced by protein deprivation. Protein restriction triggers the degradation of phosphatidylethanolamine (PE), releasing the nitrogen-containing headgroup, ethanolamine, which subsequently serves a dual function. Ethanolamine stimulates protein intake by activating protein hunger neurons and, remarkably, along with phosphoethanolamine, donates nitrogen for amino acid biosynthesis. This nitrogen transfer is mediated by the microbiome and ethanolamine-phosphate phospho-lyase (ETNPPL) pathway in fly cells. Together, our findings identify phospholipids as a hidden nitrogen reservoir and reveal an unrecognized metabolic plasticity that reallocates nitrogen to maintain protein homeostasis.

## Introduction

Macronutrients, including carbohydrates, fats, and proteins, form the essential foundation for the survival and well-being of all living animals. In preparation for future periods of food scarcity, animals store extra carbohydrates as glycogen in the liver and muscle, and surplus fats as triglycerides in adipose tissue. Excess proteins, in contrast, are not stored in a dedicated reservoir. Instead, surplus amino acids, the building blocks of protein, are catabolized and excreted (Rodwell et al 2003). While muscles can store some amino acids, this capacity is limited compared to the massive storage potential of adipose tissue for fats (Simpson & Raubenbeimer 2012). Compounding this issue, animals are also incapable of synthesizing essential amino acids and, therefore, must obtain them from their diet. The lack of a dedicated storage site, in conjunction with the challenge of de novo synthesis, necessitates a continual dietary intake of protein to fulfill physiological demands, especially during anabolic states such as pregnancy or infancy. Protein deficiency can have detrimental effects on health and survival, leading to conditions such as muscle wasting, compromised immune function, and reproductive issues (Marty et al 2017, Muller & Krawinkel 2005).

Despite lacking protein storage capacity, protein-specific hunger, which motivates animals to seek and consume a protein-enriched diet across various species, typically emerges only after prolonged protein deprivation lasting days or longer (Griffioen-Roose et al 2012, Peregoy et al 1972, Ribeiro & Dickson 2010). To cope with mild protein insufficiency and the concomitant nitrogen limitation, one strategy humans employ is reducing nitrogen excretion by decreasing urea production and increasing nitrogen salvaging from urea (Danielsen & Jackson 1992). Hibernating squirrels adopt a similar approach to preserve muscle mass (Regan et al 2022). However, this alone is insufficient to fully restore protein homeostasis (Danielsen & Jackson 1992). Could animals possess an as-yet-undiscovered strategy to derive protein from alternative sources before they are compelled to seek protein-enriched diets? Could there be a “hidden” nitrogen reservoir in the body?

Recent studies in *Drosophila* fruit flies have advanced our understanding of the cellular and molecular substrates that mediate the homeostatic regulation of protein feeding (Kim et al 2021, Leitao-Goncalves et al 2017, Li et al 2024, Munch et al 2022, Ro et al 2016, Steck et al 2018, Sun et al 2017, Yang et al 2018, Yoshinari et al 2024). Among these, a group of dopaminergic neurons projecting to the wedge neuropil, DA-WED, are critical for protein-specific hunger, with their activation driving persistent protein consumption (Liu et al 2017), and their resting membrane potential programming the protein intake setpoint (Wu et al 2024). The role of the gut-brain axis in regulating protein hunger is also increasingly recognized. Notably, gut microbes have been reported to supply essential amino acids to both insect and human hosts, thereby suppressing protein-specific hunger (Douglas & Prosser 1992, Henriques et al 2020, Kim et al 2021, Leitao-Goncalves et al 2017, Metges 2000). However, it remains debated whether essential amino acids from the microbiome suppress protein hunger by directly augmenting the host amino acid pool (Henriques et al 2020).

Here, using *Drosophila*, we discovered a previously unknown nitrogen flow from phospholipids to amino acids, which is elevated under protein starvation conditions.

Ethanolamine, the nitrogen-containing headgroup of the phospholipid phosphatidylethanolamine (PE), when released, has two key roles in defending protein homeostasis. First, ethanolamine signals protein scarcity to the brain by stimulating the DA-WED protein hunger neurons, driving an increased preference for protein-enriched food. Secondly, and remarkably, ethanolamine, along with its phosphorylated form, phosphoethanolamine, acts as a nitrogen carrier that circulates throughout the body. Facilitated by the microbiome and the ethanolamine-phosphate phospho-lyase (ETNPPL) pathway in fly cells, nitrogen from ethanolamine and phosphoethanolamine is extracted and incorporated into amino acids for protein biosynthesis.

Blocking nitrogen utilization from ethanolamine and phosphoethanolamine significantly impairs female fertility. Together, our findings identify PE as a “hidden” nitrogen reservoir in the fly. In the absence of sufficient dietary protein, nitrogen is mobilized from this reservoir to compensate for amino acid loss, with protein-specific hunger subsequently triggered to promote dietary intake to restore protein homeostasis.

## Results

### Protein starvation raises circulating ethanolamine and phosphoethanolamine

To monitor the progression of protein-specific hunger in individual flies in a high throughput manner, we developed a single fly two-choice plate feeding assay (**Figure 1A**). In this assay, two 96-well plates, each containing one type of dye-colored food, are aligned, to form 96 independent two-choice feeding chambers. Ingested dyes analyzed at their corresponding absorbances are used to calculate the intake amount for each food and the food preference index. To validate the plate assay, we examined food preference in wild-type flies subjected to protein deprivation on a sugar-only diet. Instead of mated females, we used male and virgin female flies, as these flies exhibit a lower protein preference at baseline. Consistent with previous observations under these conditions with a group of flies (Dus et al 2011, Liu et al 2017), extended protein starvation gradually increased protein preference in both virgin females and males, while amino acid supplementation suppressed this increase (**Figures S1A and S1B**). To track the progression of protein hunger without interference from deficiencies of other nutrients in the sugar-only diet, we switched to a holidic diet (Piper et al 2014) for protein deprivation. By eliminating only amino acid mix from the complete holidic diet, the plate assay again detected a gradual increase in protein preference among both virgin females (**Figures 1B and 1C**) and males (**Figure 1D**), with a peak observed around days 5 and 7, when most flies showed a preference for protein.

**Figure 1.**
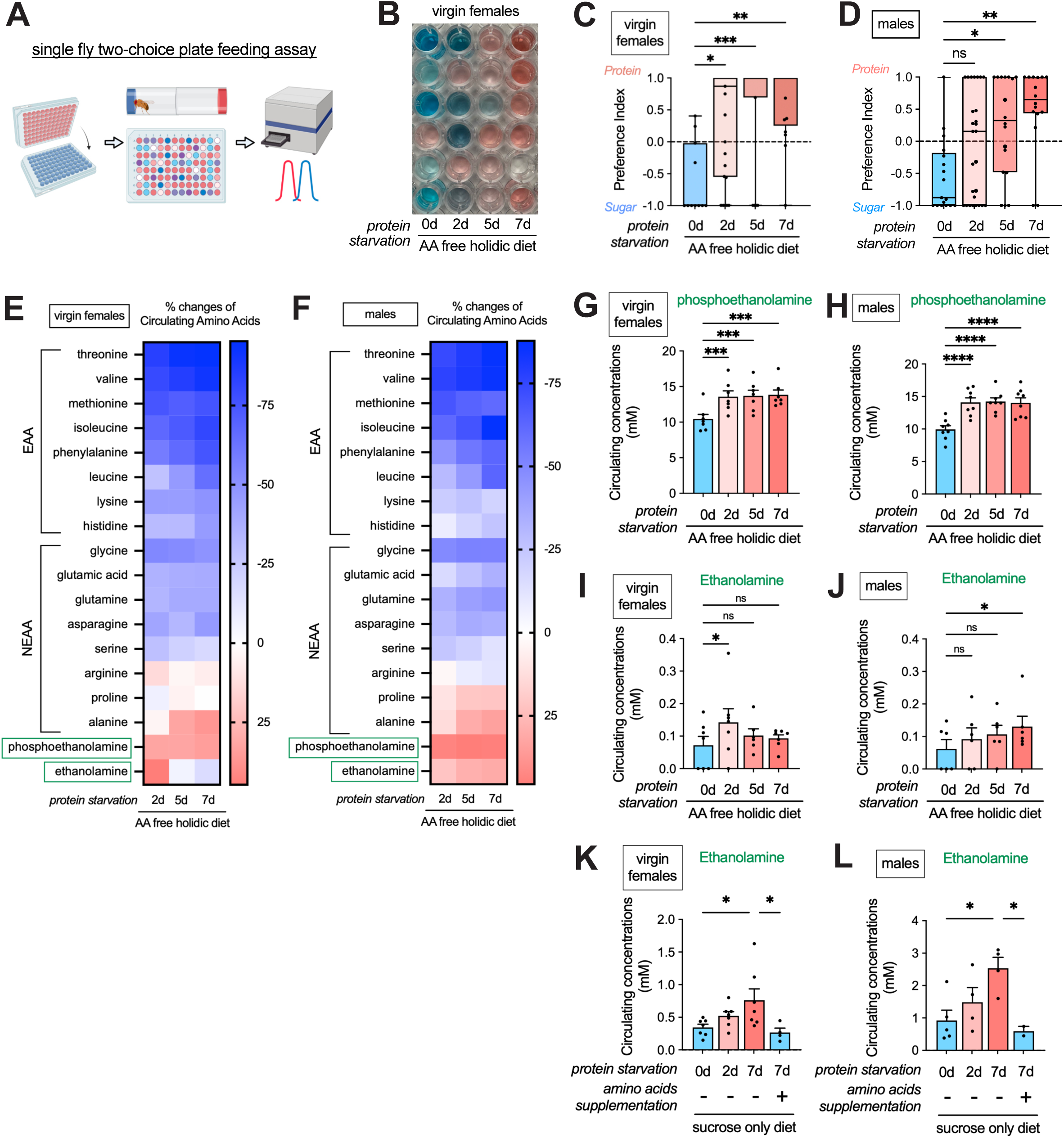
Protein starvation raises circulating ethanolamine and phosphoethanolamine. (A) Schematic of single fly two-choice plate feeding assay. Two 96-well plates are each filled with a single type of dye-colored food. Flies are introduced into one plate following cold anesthetization, and the second plate is aligned to create 96 independent feeding chambers. After a 2-hour feeding period, flies are sacrificed by freezing, and dye within their abdomens is extracted by smashing the flies and dissolving the dye in water. Color intensity and composition are then analyzed using a plate reader, measuring at 628nm for blue dye and 505nm for red dye. The absorbances are used to calculate the amount of food intake and food preferences. (B) A representative image of the two-choice feeding assay, showing dye extraction from individual age-matched wild-type *iso31* virgin female flies subjected to protein starvation on an amino acid-free holidic diet for 0, 2, 5, and 7 days. (**C-D**) Protein preference index in age-matched wild-type *iso31* virgin female (**C**) and male (**D**) flies subjected to protein starvation on an amino acid-free holidic diet for 0, 2, 5, and 7 days. n=11-25 in **C**, n=17-27 in **D**. (**E-F**) % changes in circulating amino acids from the hemolymph of age-matched wild-type virgin female (**E**) and male (**F**) flies following protein starvation on an amino acid-free holidic diet for 2, 5, and 7 days, compared to control flies without protein starvation. n=7 replicates in **E** and 8 in **F**. Each replicate contains hemolymph collected from at least 200 flies per condition. (**G-J**) The same dataset as in **E-F**, showing the concentrations of circulating phosphoethanolamine (**G-H**) and ethanolamine (**I-J**) in the hemolymph of age-matched wild-type virgin female and male flies subjected to protein starvation on an amino acid-free holidic diet for 0, 2, 5, and 7 days. n=7 replicates in **G** and **H**, n=8 replicates in **I**, and n=6 replicates in **J** due to the elimination of datasets containing undetectable ethanolamine. Each replicate contains hemolymph collected from at least 200 flies per condition. (**K-L**) Concentrations of circulating ethanolamine in the hemolymph of age-matched wild-type virgin female (**K**) and male (**L**) flies subjected to protein starvation on a sucrose-only diet for 0, 2, and 7 days with or without amino acids supplementation. n=7 replicates in **K**, n=2-5 replicates in **L**. Each replicate contains hemolymph collected from at least 200 flies per condition. Kruskal-Wallis tests were performed with Dunn’s multiple comparisons test for Figures 1C and **1D**. Repeated measures one-way ANOVA with Dunnett’s multiple comparisons test was used to analyze data in Figures 1G**-1L**. In this and the subsequent figures, for normally distributed datasets, bar plots with mean ± SEM are shown, with scattered dots representing individual data points. For non-normally distributed datasets, box plots are shown with scattered dots to display all raw data points. The line inside the box indicates the median. The top and bottom of the box represent the 75th and 25th percentiles, respectively. The whiskers represent the minimum and maximum values. “*”, “**”, “***”, “****”, and “ns” denote P<0.05, P<0.01, P<0.001, P<0.0001, and not significant, respectively. See also **Figure S1**.

Using the same protein starvation scheme, we next sought to assess the metabolic changes associated with the lack of dietary protein intake, starting by measuring the circulating levels of amino acids in hemolymph, which is analogous to blood in vertebrates. Interestingly, in both virgin female and male flies, most detectable essential amino acids showed a marked decrease within two days of protein starvation and remained low with prolonged deprivation (**Figures 1E and 1F, S1C and S1D**). In contrast, non-essential amino acids exhibited either a milder decrease or, in some cases, a slight increase, as seen with arginine, proline, and alanine. Notably, circulating levels of phosphoethanolamine—the phosphorylated form of the amino alcohol ethanolamine and one of the most abundant compounds in our amino acid analysis—showed a robust increase after two days of protein deprivation, remaining elevated with continued starvation in both virgin female and male flies (**Figures 1E-1H**). Ethanolamine, on the other hand, was present at much lower baseline concentrations and was often undetectable (**Figures 1I and 1J**). Following protein starvation, ethanolamine levels showed a mild elevation, becoming more consistently detectable and significantly higher than controls on day 2 in virgin females and day 7 in males (**Figures 1I and 1J**).

Due to the low baseline levels of ethanolamine on the holidic diet, we validated the data using the earlier protein deprivation scheme, with regular food as the control and sucrose-only diet for deprivation (**Figures S1A and S1B**). In both virgin females and males, protein deprivation on the sucrose-only diet similarly triggered a decrease in most amino acids, which was largely rescued by supplementation with an amino acid mix (**Figures S1E and S1F**). Under these conditions, circulating ethanolamine levels were higher at baseline compared to the holidic diet, allowing for consistent detection. Ethanolamine levels gradually increased with prolonged protein starvation and were restored to baseline levels by amino acid supplementation (**Figures 1K and 1L**), tightly correlating with behavioral changes in protein preference (**Figures S1A and S1B**). Meanwhile, circulating phosphoethanolamine levels followed a similar pattern in virgin females (**Figure S1E**) but did not increase in males (**Figure S1F**) during protein starvation.

To determine whether these changes are specifically associated with protein starvation, we next examined the effects of complete food deprivation on circulating amino acid levels. Total food starvation for 24 hours did not consistently decrease circulating amino acid levels. Instead, some essential amino acids, including threonine, lysine, and histidine, showed a slight increase (**Figure S1G**). While circulating phosphoethanolamine levels increased under total food starvation, mirroring the response to protein starvation, ethanolamine was markedly downregulated in both virgin females and males following complete food starvation (**Figure S1G**). These findings indicate that the increase in ethanolamine, but not phosphoethanolamine, is specifically associated with protein starvation rather than general hunger. Collectively, our data reveal distinctive circulating metabolic changes induced by protein starvation, including a pronounced decrease in essential amino acids and a substantial increase in ethanolamine and phosphoethanolamine, with the increase in ethanolamine being specific to protein starvation across sexes and dietary conditions.

### Ethanolamine activates the DA-WED protein hunger neurons

Ethanolamine and phosphoethanolamine are well recognized as the head group of the phospholipid Phosphatidylethanolamine (PE), a major component of the eukaryotic cell membrane (**Figure S2A**). Emerging data suggest that ethanolamine may exert its own functions, independent of phospholipids, in cell signaling and synaptic transmission (Gwanyanya et al 2022, Kendall et al 2012). Does increased ethanolamine and phosphoethanolamine in circulation play an active role in protein-specific hunger, or are they only byproducts of metabolic changes? To address this, we artificially increased ethanolamine and phosphoethanolamine levels via dietary supplementation and assessed the behavioral consequences on food preference. Under protein-satiated conditions, ethanolamine supplementation increased protein preference in a dose- dependent manner, in both virgin female (**Figures 2A and 2B**) and male (**Figure S2B**) flies. In contrast, supplementation of phosphoethanolamine up to 100 mM did not induce protein preference in male flies (**Figure S2C**). In virgin female flies, phosphoethanolamine supplementation at 25 mM showed a trend toward increasing protein preferences, though this effect was not observed at lower or higher concentrations (**Figure S2D**). These data indicate that ethanolamine, not phosphoethanolamine, can actively induce protein-specific hunger.

**Figure 2.**
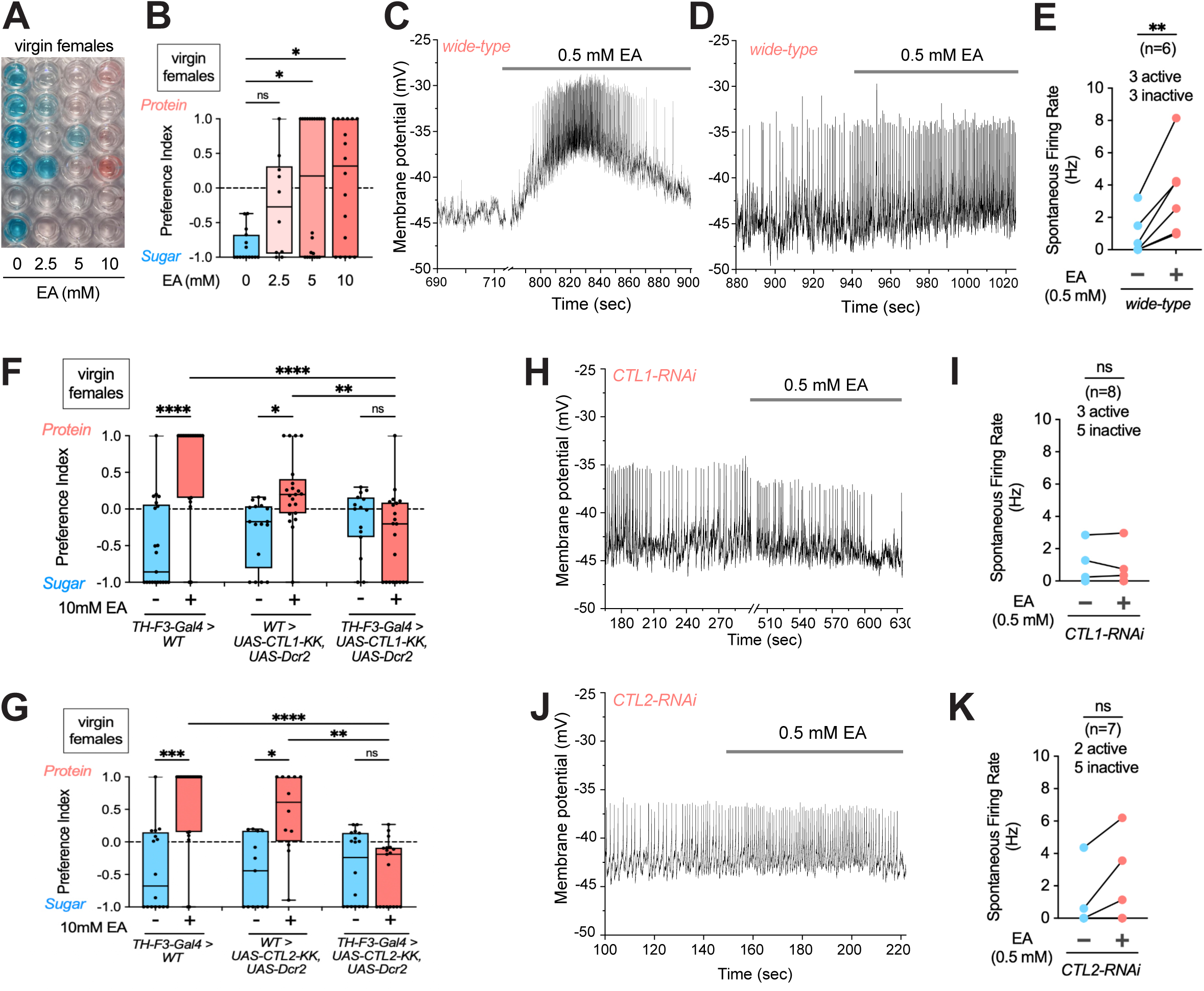
Ethanolamine activates the DA-WED protein hunger neurons. (**A-B**) Representative image of a two-choice plate (**A**) and quantified protein preference index (**B**) for age-matched wild-type *iso31* virgin female flies, fed on complete holidic diets supplemented with 0mM, 2.5mM, 5mM, or 10mM ethanolamine for 3 days prior to two-choice plate feeding assay. Flies were first entrained on a complete holidic diet for 2 days before being transferred to ethanolamine-supplemented holidic diets. n=10-22. (**C-E**) Representative traces (**C-D**) and action potential firing rate (**E**) for patch clamp recordings of DA-WED neurons from *TH-F3-Gal4>UAS-mCD8-GFP* virgin female flies before and after bath application of 0.5mM ethanolamine. n=6. (**F**) Protein preference index in virgin female flies of controls (*TH-F3-Gal4>WT* and *WT>UAS-CTL1-KK-RNAi, UAS-Dcr2*) or CTL1 knockdown in DA-WED neurons (*TH-F3-Gal4>UAS-CTL1-KK-RNAi, UAS-Dcr2*), with or without supplementation of 10mM ethanolamine in a complete holidic diet for 3 days. n=15-21. (**G**) Protein preference index in virgin female flies of controls (*TH-F3-Gal4>WT* and *WT>UAS-CTL2-KK-RNAi, UAS-Dcr2*) or CTL2 knockdown in DA-WED neurons (*TH-F3-Gal4>UAS-CTL2-KK-RNAi, UAS-Dcr2*), with or without supplementation of 10mM ethanolamine in a complete holidic diet for 3 days. n=12-21. (**H-I**) Representative traces (**H**) and action potential firing rate (**I**) for patch clamp recordings of DA-WED neurons from *TH-F3-Gal4>UAS-mCD8-GFP, UAS-CTL1-RNAi, UAS-Dcr2* virgin female flies before and after bath application of 0.5mM ethanolamine. n=8. (**J-K**) Representative traces (**J**) and action potential firing rate (**K**) for patch clamp recordings of DA-WED neurons from *TH-F3-Gal4>UAS-mCD8-GFP, UAS-CTL2-RNAi, UAS-Dcr2* virgin female flies before and after bath application of 0.5mM ethanolamine. n=7. Kruskal-Wallis tests with Dunn’s multiple comparisons test were performed for Figures 2B, **2F,** and **2G**. Wilcoxon matched-pairs test was used to analyze data in Figures 2E, **2I,** and **2K**. See also **Figure S2**.

How does ethanolamine trigger protein-specific hunger? There is evidence suggesting the role of ethanolamine as a neuromodulator in various species (Gwanyanya et al 2022). DA-WED is a group of dopamine neurons in the central brain of *Drosophila* that are activated by protein starvation and subsequently stimulate protein consumption (Liu et al 2017). We thus asked whether ethanolamine modulates the activities of DA-WED neurons, using patch-clamp recordings. We prioritized virgin females due to their larger cell body size, resulting in a higher success rate of patch-clamp. At baseline, a subset of DA-WED neurons (3 out of 6 neurons recorded) do not exhibit spontaneous action potential firing. Bath application of 0.5 mM ethanolamine induced firing in all of these previously inactive neurons (**Figures 2C and 2E**). In neurons that displayed spontaneous firing, ethanolamine also significantly increased their firing rate (**Figures 2D and 2E**). On the other hand, bath application of phosphoethanolamine at 0.5mM, or as low as 0.1mM, disrupts giga seal and prevents signal acquisition in patch-clamp recordings. These data support the notion that ethanolamine functions as a neuromodulator, exciting the protein hunger neurons to promote protein feeding.

We next sought to elucidate the molecular pathways mediating the neuromodulatory effect of ethanolamine in DA-WED neurons. A recent in vitro study established the role of choline transporter-like proteins-1 and -2 (CTL1 and CTL2), the solute carriers 44A (SLC44A) family members, in transporting ethanolamine at the plasma and mitochondrial membranes in human fibroblast cells (Taylor et al 2021). Fly genome also encodes two choline transporter-like proteins, CTL1 and CTL2, the physiological function of which remains elusive. We hypothesize that fly CTL1 and CTL2 facilitate the transport of ethanolamine from extracellular space to DA-WED neurons, thereby mediating the activation of these cells upon protein starvation. To test this, we suppressed the expression of CTL1/2 in DA-WED neurons and assessed the impact on ethanolamine-induced protein hunger. CTL1 Knockdown in DA-WED neurons effectively blocked the protein preference triggered by ethanolamine supplementation (**Figure 2F**). Similar results were obtained using an RNAi line targeting CTL2 (**Figure 2G**). Consistent with the behavioral data, in flies carrying CTL1 RNAi, bath application of ethanolamine did not increase the firing rate of either active or inactive DA-WED neurons (**Figures 2H and 2I**). In flies carrying CTL2 RNAi, ethanolamine failed to induce firing in 4 out of 5 previously inactive DA- WED neurons, and modestly increased the firing rate in 2 active neurons (**Figure 2J**), although the changes were not statistically significant (**Figures 2K**). These data suggest that both CTL1 and CTL2 are required for ethanolamine-mediated activation of DA-WED neurons. Together, our findings highlight the role of ethanolamine as a messenger that signals protein deficiency to protein hunger neurons, and identify CTL1 and CTL2 as key transporters that mediate its neuromodulatory effects.

### Protein starvation triggers the breakdown of phospholipid PE

What are the sources of increased ethanolamine in circulation and the brain following protein starvation? Since animals do not synthesize ethanolamine de novo, they mostly source it from their diet, either as free ethanolamine or in the form of PE, which releases ethanolamine upon degradation (Gibellini & Smith 2010, Lee et al 2019) (**Figure S2A**). Ethanolamine can also originate from the degradation of other molecules, such as sphingosine phosphate and endocannabinoid anandamide, but the abundance of these alternative sources is extremely low compared to PE (Hannun et al 2001, Matas et al 2007). Given that PE is not provided in the holidic diet, we hypothesized that the degradation of internal PE contributes to the elevated ethanolamine levels following protein starvation. This predicts that suppressing PE breakdown would prevent the increase in ethanolamine levels and, consequently, reduce protein preference. As Phospholipase D (PLD) cleaves the headgroup from phospholipids in both mammals and flies (**Figure 3A**) (LaLonde et al 2005, Panda et al 2018, Sandoval 2012), we evaluated the behavioral impact in a *pld* mutant strain, *pld3.1* (Thakur et al 2016). The *pld3.1* mutant exhibited a significantly delayed response to protein starvation compared to wild-type, with no increase in protein preference until 7 days after protein deprivation (**Figure 3B**). This result suggests that elevated ethanolamine, potentially generated through PLD-mediated PE degradation, is essential for signaling protein-specific hunger in a timely manner.

**Figure 3.**
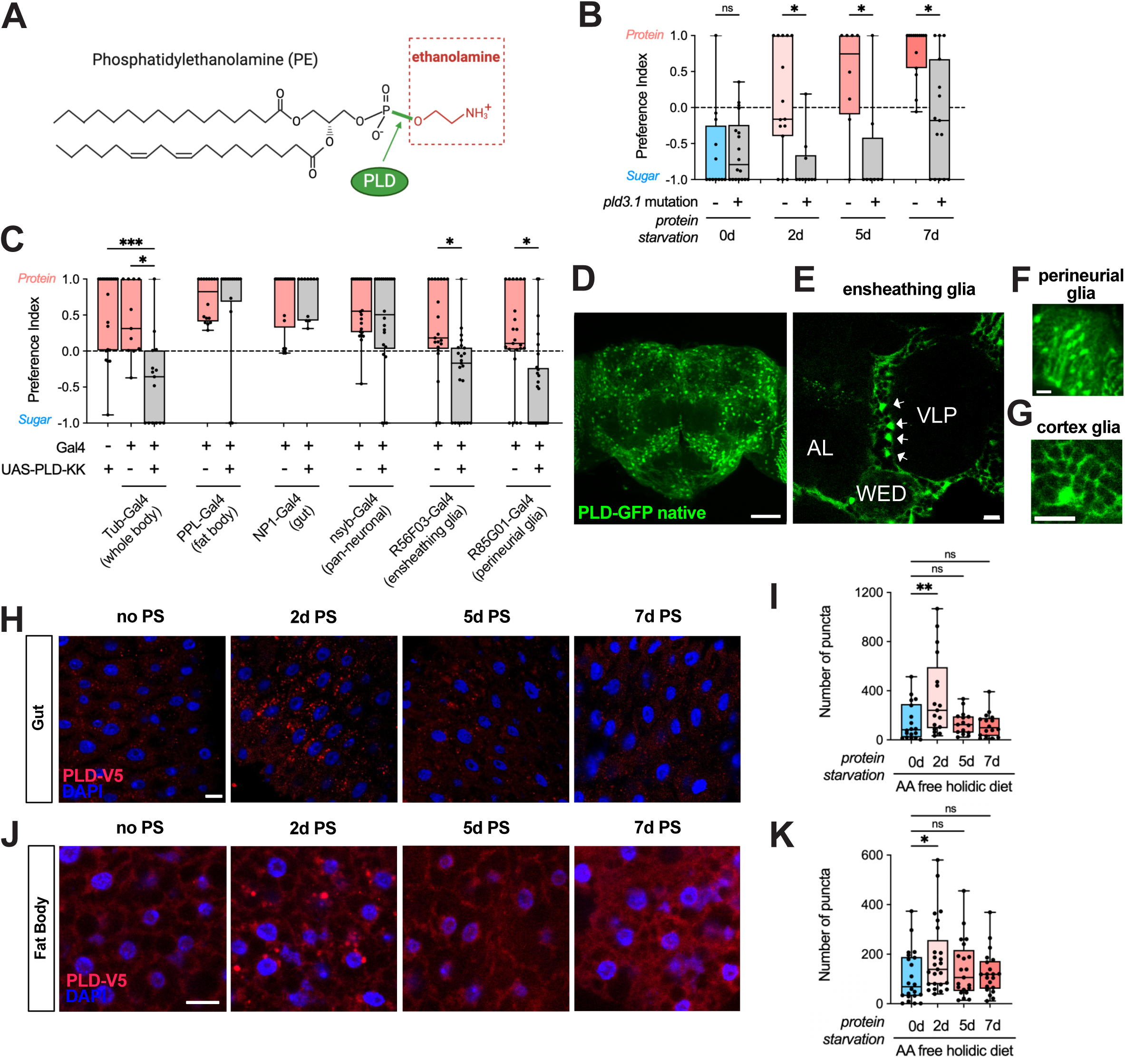
Protein starvation triggers the breakdown of phospholipid PE. (A) Structure of phosphatidylethanolamine (PE), with the headgroup ethanolamine highlighted in red and the phospholipase D (PLD) cleavage site highlighted in green. (B) Protein preference index in virgin female flies of wild-type or *pld3.1* mutant flies subjected to protein starvation on an amino acid-free holidic diet for 0, 2, 5, and 7 days. n=8-18. (C) Protein preference index in virgin female flies of controls (*Gal4>WT* and *WT>UAS-PLD-KK RNAi*) or tissue-specific PLD knockdown in the whole body (*Tub-Gal4>UAS-PLD-KK RNAi*), fat body (*PPL-Gal4>UAS-PLD-KK RNAi*), gut (*NP1-Gal4>UAS-PLD-KK RNAi*), all neurons (*nsyb-Gal4>UAS-PLD-KK RNAi*), ensheathing glia (*R56F03-Gal4>UAS-PLD-KK RNAi*), or perineurial glia (*R85G01-Gal4>UAS-PLD-KK RNAi*), following protein starvation on an amino acid-free holidic diet for 5 days. n=11-28. (**D-G**) Whole-mount brain images of the PLD-GFP-V5 strain (VK00033) from the fTRG library, showing native GFP fluorescence in a maximum whole-brain projection (**D**), and single z-stack images highlighting ensheathing glia (**E**), perineurial glia (**F**), and cortex glia (**G**). In **E**, arrows indicate ensheathing glia nuclei located at the boundaries between neuropils: AL (antennal lob), VLP (ventral lateral protocerebrum), and WED (wedge). Scale bar, 50 μm in **D**, 10 μm in **E**-**G**. (**H-I**) Representative images (**H**) and number quantification (**I**) of PLD-GFP-V5 puncta in the enterocytes of the R3 gut sections from age-matched virgin female flies subjected to protein starvation (PS) on an amino acid-free holidic diet for 0, 2, 5, and 7 days, Samples were immunostained with antibodies against V5 (red) and counterstained with nuclear marker DAPI (blue). n=15-18. Scale bar, 10 μm. (**J-K**) Representative images (**J**) and number quantification (**K**) of PLD-GFP-V5 puncta in fat body cells from age-matched virgin female flies subjected to protein starvation (PS) on an amino acid-free holidic diet for 0, 2, 5, and 7 days, Samples were immunostained with antibodies against V5 (red) and counterstained with nuclear marker DAPI (blue). n=22-27. Scale bar, 10 μm. Kruskal-Wallis tests with Dunn’s multiple comparisons test were performed for Figures 3B**, 3C, 3I, 3K.** See also **Figure S3**.

To determine the specific tissues where PLD functions to cleave PE and release ethanolamine in response to protein starvation, we performed tissue-specific PLD knockdown using RNAi. As expected, PLD knockdown in all cells with tubulin-Gal4 phenocopied *pld3.1* mutant, suppressing protein preference in protein-deprived animals (**Figure 3C**). However, knockdown of PLD in the fat body (*PPL-Gal4*), gut (*NP1-Gal4*), or neurons (*nsyb-Gal4*) did not significantly impact protein preference. The single-cell transcriptomic dataset revealed that PLD expression in the brain is enriched in glial cells, specifically ensheathing glia and perineurial glia (**Figure S3A**) (Dopp et al 2024). This glial enrichment was also observed in body samples containing the ventral nerve cord (Li et al 2022). To test the functional significance of the glial localization, we knocked down PLD using driver lines specific to neuropile ensheathing glia (*R56F03-Gal4*) or perineurial glia (*R85G01-Gal4*) (Kremer et al 2017). Notably, both driver lines were sufficient to suppress protein preference when combined with PLD RNAi (**Figure 3C**), suggesting that PLD is required “locally” in the brain glial cells to generate ethanolamine and drive protein preference in response to protein deprivation.

Using a C-terminally tagged PLD strain from the fTRG library (Sarov et al 2016), we examined the expression of PLD-GFP-V5 chimeric protein in the brain. Native GFP fluorescence was readily detected in the ensheathing glia, characterized by nuclei positioned along neuropil boundaries and processes extending to wrap around the neuropil edges (**Figures 3D and 3E**).

Although the signal from perineurial glia was largely masked, we occasionally identified these cells at the surface area as narrow, oblong structures (**Figure 3F**). Additionally, we observed cortex glia in the cortical regions, where they envelop neuronal cell bodies individually, forming cuboid-shaped compartments (**Figure 3G**). Although native GFP fluorescence was too weak to be observed in the gut and fat body, we were able to detect the expression of PLD-GFP-V5 in these tissues using a V5 antibody. Notably, while PLD knockdown in gut and fat body cells did not affect protein preference, protein starvation for two days significantly increased the PLD-V5 signal, marked by a rise in both puncta number and size (**Figures 3H-3K and Figures S3B-S3C**). However, prolonged protein starvation did not further elevate PLD-V5 levels. This pattern parallels the increase in circulating ethanolamine in the hemolymph, which also peaks at day 2 in virgin females (**Figure 1I**). These findings suggest the upregulation of PLD expression in peripheral tissues, including the gut and fat body, may contribute to the observed rise in circulating ethanolamine levels under protein starvation. Together, our results support the notion that protein starvation promotes PE breakdown, thereby releasing its headgroup, ethanolamine.

### Nitrogen flow from ethanolamine and phosphoethanolamine to amino acids

Our data so far suggest that protein starvation increases the release of ethanolamine from PE in both the brain and the peripheral tissues. However, ethanolamine produced in brain glial cells—not the peripheral tissues—is required to induce protein preference. This raises an important question: what role do the increased levels of circulating ethanolamine and phosphoethanolamine play? Since ethanolamine and phosphoethanolamine contain nitrogen, we wondered if this nitrogen could be extracted and utilized for amino acid biosynthesis. Although unconventional, we set out to explore this possibility by labeling the nitrogen in ethanolamine with ^15^N, a stable isotope of the naturally prevalent ^14^N. By supplementing the fly diet with ^15^N-EA, we investigated whether ^15^N was incorporated into amino acids, using compound-specific stable isotope analysis of nitrogen (**Figure 4A**). In the absence of ^15^N-EA supplementation, ^15^N-amino acid relative to total amino acids remained near zero, reflecting the extremely low levels of naturally occurring ^15^N-amino acids (**Figure 4B**). After 5 days of incubation on a ^15^N-EA supplemented diet, we observed a drastic increase in the fraction of ^15^N-amino acid for all detected essential and non-essential amino acids, in a ^15^N-EA dose-dependent manner (**Figure 4B**). This confirms that the nitrogen from supplemented ethanolamine was successfully extracted and utilized for de novo amino acid biosynthesis.

**Figure 4.**
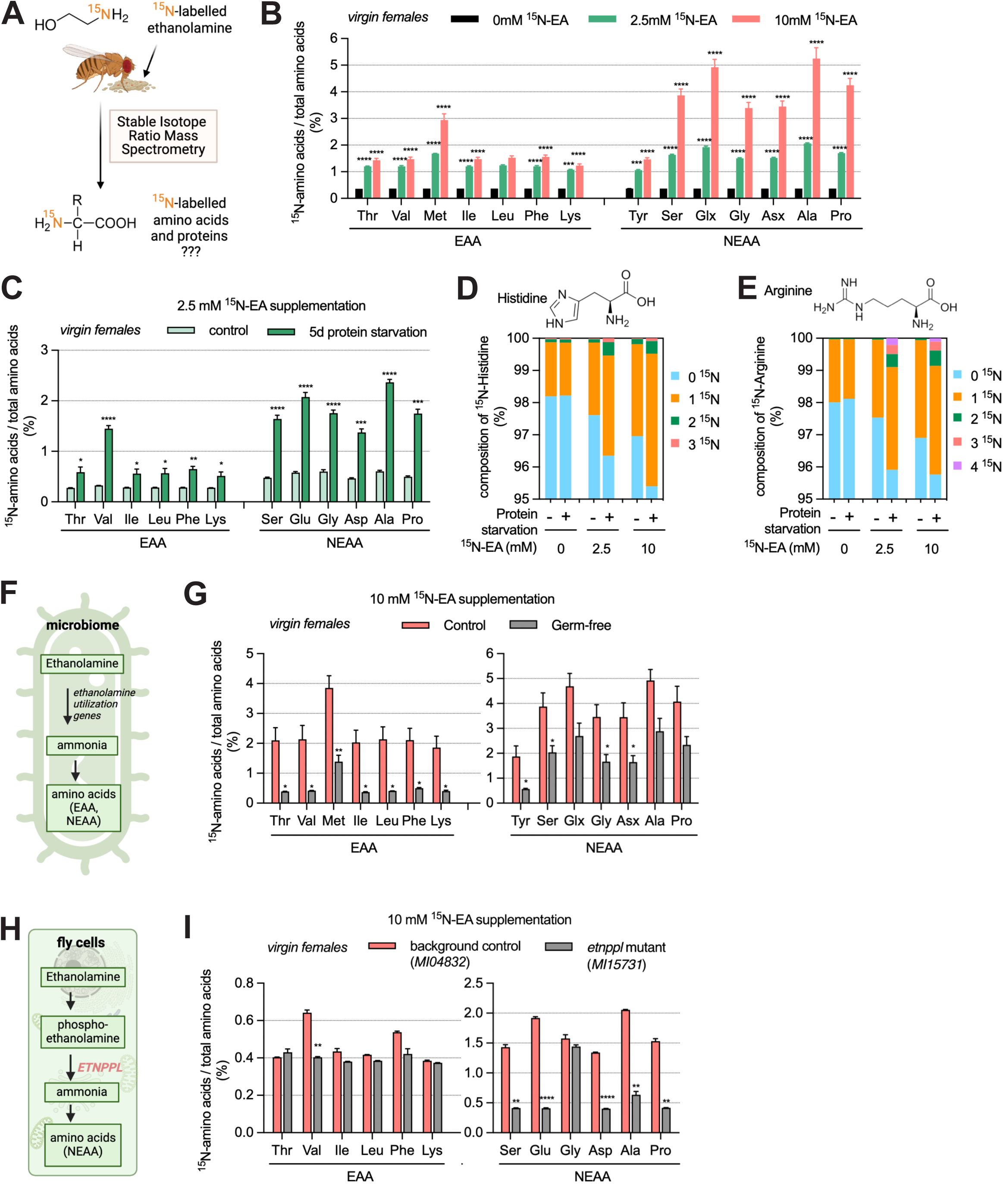
Nitrogen flow from ethanolamine and phosphoethanolamine to amino acids. (A) Schematic of the isotope tracing of nitrogen in ethanolamine. The nitrogen in ethanolamine is labeled with ^15^N, a stable isotope of the naturally prevalent ^14^N. ^15^N-ethanolamine (^15^N-EA) is then supplemented into the holidic fly diet. Following 5 days of feeding on the ^15^N-EA supplemented diet, whole-body fly samples are subjected to stable isotope analyses to assess the incorporation of ^15^N into amino acids. (B) Percentage of ^15^N-amino acid relative to total amino acids (including both free-amino acids and those incorporated into proteins) in age-matched wild-type *iso31* virgin female flies fed on holidic diets supplemented with 0mM, 2.5mM, and 10mM ^15^N-EA. n= 3 replicates, with each replicate containing 20-30 flies per condition. Glx represents the combined measurement of glutamine (Gln) and glutamate (Glu), while Asx represents the combined measurement of aspartate (Asp) and asparagine (Asn), due to the inability to separate them under this experimental setting. (C) Percentage of ^15^N-amino acid relative to total amino acids in age-matched wild-type *iso31* virgin female flies fed for 5 days on 2.5mM ^15^N-EA supplemented complete holidic diets (control) or amino acid-free holidic diet (5d protein starvation). n=3 replicates, with each replicate containing 20-30 flies per condition. (**D-E**) Percentage for each ^15^N-labeled heavy form, containing different numbers of ^15^N, relative to total Histidine (**D**) and Arginine (**E**) in age-matched wild-type *iso31* virgin female flies fed on holidic diets supplemented with 0mM, 2.5mM, and 10mM ^15^N-EA, with or without protein starvation. n= 3 replicates, with each replicate containing 20-30 flies per condition. (F) Schematic of the process where bacteria convert ethanolamine into ammonia and acetate, facilitated by ethanolamine utilization (EUT) genes. The ammonia is subsequently being incorporated into amino acid biosynthesis. (G) Percentage of ^15^N-amino acids relative to total amino acids in age-matched control and germ-free virgin female flies fed for 5 days on 10mM ^15^N-EA supplemented complete holidic diets. n= 3 replicates, with each replicate containing 20-30 flies per condition. (H) Schematic of the pathway in host cells where phosphoethanolamine is degraded by the enzyme ethanolamine-phosphate phospho-lyase (ETNPPL), releasing ammonia, which is subsequently incorporated into the biosynthesis of non-essential amino acids. (I) Percentage of ^15^N-amino acids relative to total amino acids in age-matched background control *MI04832* (BDSC_38587) and *etnppl* mutant *MI15731*(BDSC_61139) virgin female flies fed for 5 days on 10mM ^15^N-EA supplemented complete holidic diets. n=2 replicates, with each replicate containing 20-30 flies per condition. Two-way ANOVA with the Geisser-Greehouse correction, followed by Tukey’s multiple comparisons test, was used to analyze the data in Figure 4B. Multiple Unpaired t-tests with Welch correction were performed for Figures 4C, **4G**, and **4I**. See also **Figure S4**.

We next tested the impact of protein starvation on the incorporation of nitrogen from ethanolamine to amino acids. In response to 5-day protein starvation, the fraction of ^15^N-amino acids substantially increased in animals fed on both 2.5mM (**Figure 4C**) and 10mM ^15^N-EA (**Figure S4A**) supplemented diet, an effect that was not observed without ^15^N-EA supplementation (**Figure S4B**). Since we had previously observed a robust decrease in circulating amino acids in protein-starved animals (**Figures 1E and 1F**), we wondered if the increased proportion of ^15^N-amino acids was simply due to an overall reduction in total amino acids following protein deprivation. To clarify this, we calculated the peak signals for both ^15^N- and ^14^N-amino acids, as an estimate of the absolute amount of these amino acids in the whole-body samples. We found that ^14^N-amino acids showed no significant decrease except for Val and Thr (**Figures S4C and S4E**), while ^15^N-amino acids, in particular non-essential amino acids, increased following protein starvation (**Figures S4D and S4F**). This confirms that the elevated proportion of ^15^N-amino acids in protein-starved animals was not due to a decrease in total amino acids. Additionally, these changes cannot be attributed to differences in food intake, as the total 5-day food consumption, measured using the lid assay (Wu et al 2024), showed no significant differences across the various feeding conditions (**Figure S4G**). Another interesting observation is that amino acids containing more than one nitrogen atom, such as histidine (3N) and arginine (4N), exhibited multiple ¹⁵N-labeled “heavy forms”. In response to protein starvation, the fraction of histidine and arginine containing 1 or 2 ¹⁵N atoms significantly increased. Furthermore, protein starvation induced the incorporation of 3 ¹⁵N atoms into histidine and 3 or 4 ¹⁵N atoms into arginine, which was not detected in non-starved flies (**Figures 4D and 4E**). This suggests that nitrogen from ethanolamine can be incorporated into various parts of the amino acid molecules.

Together, these findings demonstrate that ethanolamine can serve as a nitrogen source for amino acid biosynthesis, and this “nitrogen flow” from ethanolamine to amino acids is amplified by dietary protein restriction.

These findings are unexpected, as eukaryotic cells are generally not considered capable of catabolizing ethanolamine and using it as a nitrogen source. In contrast, certain bacteria—particularly pathogens carrying ethanolamine utilization (EUT) genes—can convert ethanolamine into ammonia and acetaldehyde, with ammonia subsequently being incorporated into amino acids (Nawrocki et al 2018) (**Figure 4F**). To ensure that the ¹⁵N-labeled amino acids we detected were from the host cells, not simply from the gut microbiome proteome, we separated fly heads and bodies for Elemental Analyzer-Isotope Ratio Mass Spectrometry (EA-IRMS), as the head contains minimal microbiota compared to the gut. The proportion of ¹⁵N incorporation was comparable between head and body samples from both control and protein-starved flies fed on a 10mM ^15^N-EA supplemented diet (**Figure S4H**), confirming that we are indeed measuring the host protein pool. Given that eukaryotic cells do not typically break down ethanolamine, we hypothesized that bacteria might facilitate the extraction of nitrogen from ethanolamine, converting it into amino acids, which are then released for utilization by the host. To test this, we examined the incorporation of ^15^N from ethanolamine to amino acids in germ-free flies.

Remarkably, in the absence of the microbiome, the levels of ¹⁵N-labeled amino acids significantly decreased for most detected amino acids (**Figure 4G**). In particular, ¹⁵N-labeled essential amino acids dropped to levels found in non-supplemented animals, except for methionine, suggesting a complete suppression of the nitrogen flow from ethanolamine to essential amino acids. Although germ-free flies consume less food compared to the controls (**Figure S4I**), this reduction alone does not appear sufficient to account for the substantial decrease in ^15^N-labeled essential amino acids observed in these animals. These findings indicate that the microbiome plays a crucial role in converting ethanolamine into essential amino acids.

The partial decrease in non-essential amino acids observed in germ-free flies (**Figure 4G**) raises the possibility that host cells may also be able to extract and utilize the nitrogen from ethanolamine, challenging the conventional views that eukaryotic cells cannot catabolize ethanolamine. Ethanolamine-phosphate phospho-lyase (ETNPPL) is a protein characterized by its ability to catalyze the degradation of phosphoethanolamine into acetaldehyde, phosphate, and ammonia with high specificity (**Figure 4H**) (Fleshood & Pitot 1970a, Schiroli et al 2013, Veiga-da-Cunha et al 2012), contributing significantly to lipid homeostasis (White et al 2021). The *Drosophila* genome encodes one ortholog of ETNPPL, CG8745, the function of which remains elusive. To investigate whether this pathway contributes to nitrogen utilization from ethanolamine in fly cells, we monitored the nitrogen flow in an *etnppl* mutant from the MiMIC library (Nagarkar-Jaiswal et al 2015). This mutant carries a MiMIC insertion (*MI15731*) in the first coding intron of *ETNPPL* gene in the same orientation, thus leading to an early stop codon and loss of function (Nagarkar-Jaiswal et al 2015). As a background control, we used another MiMIC insertion from the same library (*MI04832*), which is located in an adjacent non-coding region.

Notably, in the *etnppl* mutant, ¹⁵N incorporation into essential amino acids remained largely unaffected, except for valine (**Figure 4I**). In contrast, *etnppl* mutation significantly impaired ¹⁵N incorporation into nearly all detected non-essential amino acids except for glycine, an effect more pronounced than in germ-free flies (**Figure 4I**). This effect is not due to reduced food intake, as the total food consumption in the mutant was slightly higher than in wild-type flies (**Figure S4J**). These results suggest that fly cells also possess a mechanism for extracting and utilizing nitrogen from ethanolamine and phosphoethanolamine, which is dependent on ETNPPL. This pathway preferentially channels nitrogen from ethanolamine into non-essential amino acids, and works together with the microbiome to mobilize nitrogen from phospholipids for amino acid biosynthesis. Together, our results reveal a previously unrecognized nitrogen flow from phospholipids to amino acids, and elucidate two independent mechanisms facilitating this nitrogen flow, one contributed by the microbiome and the other by fly cells. While it is generally believed that there is no dedicated nitrogen storage site, our findings indicate that phospholipids may function as a previously unrecognized nitrogen reservoir, contributing to the maintenance of protein homeostasis.

### Impaired nitrogen utilization from phospholipids compromises female fertility

We next investigated the physiological impact of blocking nitrogen utilization from the phospholipid reservoir. Egg-laying in female flies is a process highly sensitive to both internal and external nutritional status, particularly protein availability (Alves et al 2022, Sang & King 1961). We thus first measured egg-laying numbers in *pld* mutants, germ-free flies, and *etnppl* mutants. Interestingly, even when maintained on a complete holidic diet, all three groups exhibited a significant reduction in daily egg production compared to controls (**Figures 5A-5C**). By day 5 after mating, the number of eggs laid dropped close to zero in these groups, while control flies continued to lay eggs at high levels. When flies were fed an amino acid-free holidic diet, control flies dramatically reduced egg-laying, almost completely halting by day 5, while mutants and germ-free flies exhibited an even more pronounced decline in egg-laying numbers (**Figures S5A-S5C**). To test whether protein level is the key factor underlying differences between *pld* mutants, germ-free flies, and *etnppl* mutants, and their controls, we supplemented the complete holidic diet with excess yeast, an ethologically relevant protein source. Yeast supplementation substantially increased egg laying across all groups and their controls (**Figures 5D-5F vs Figures 5A-5C**). More importantly, with excess protein, the differences in egg-laying between *pld3.1* or *etnppl* mutants and their controls were eliminated (**Figures 5D and 5F**), although germ-free flies continued to lay significantly fewer eggs than controls (**Figure 5E**).

**Figure 5.**
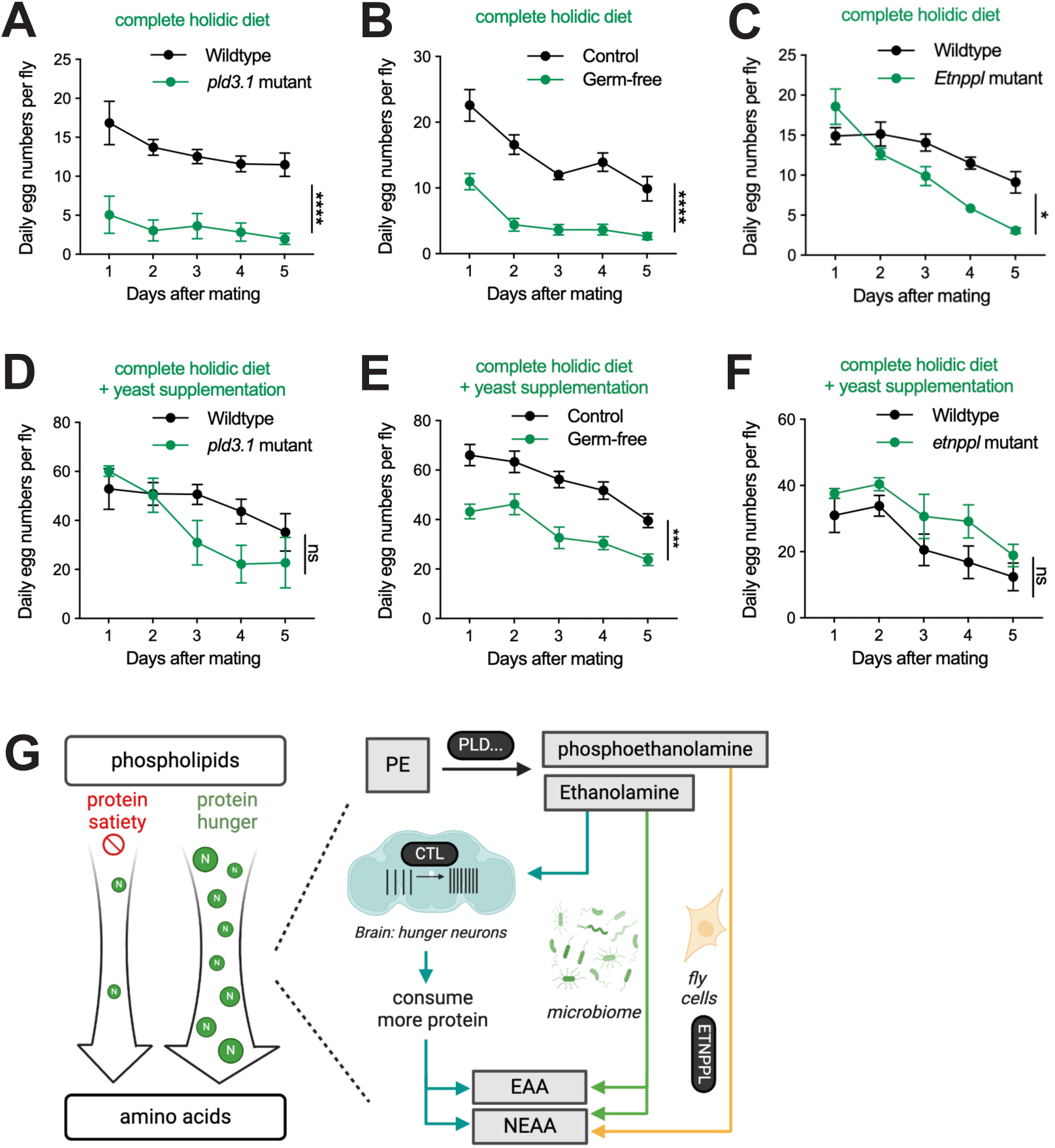
Impaired nitrogen utilization from phospholipids compromises female fertility. (**A-C**) Daily egg-laying numbers per fly in age-matched mated female flies of wild-type and *pld 3.1* mutants (**A**), control and germ-free flies (**B**), background control *MI04832* (BDSC_38587) and *etnppl* mutant *MI15731*(BDSC_61139) (**C**) on a complete holidic diet. n=8-12 replicates, with each replicate consisting of 5-6 virgin female flies mixed with 2 wild-type males. Flies were allowed to mate for 48 hours on a standard molasses diet before females were transferred to a complete holidic diet. Egg numbers were counted every 24 hours after transferring female flies into a new vial with a complete holidic diet. (**D-F**) Daily egg-laying numbers per fly in age-matched mated female flies of wild-type and *pld 3.1* mutant (**D**), control and germ-free flies (**E**), background control *MI04832* (BDSC_38587) and *etnppl* mutant *MI15731*(BDSC_61139) (**F**) on a complete holidic diet sprinkled with yeast pellets. n=3-8 replicates, with each replicate consisting of 5-6 virgin female flies mixed with 2 wild-type males per condition. Flies were allowed to mate for 48 hours on a standard molasses diet sprinkled with yeast pellets before females were transferred to a complete holidic diet sprinkled with yeast pellets. Egg numbers were counted every 24 hours after transferring female flies into a new vial with a yeast-sprinkled complete holidic diet. (**G**) Model. A nitrogen flow from phospholipids to amino acids is enhanced under protein starvation conditions. When dietary protein is insufficient, upregulated phospholipase D (PLD) and potentially other lipases promote the degradation of phospholipids, primarily phosphatidylethanolamine (PE), releasing ethanolamine and phosphoethanolamine. Ethanolamine is transported into DA-WED neurons via choline transporter-like (CTL) proteins, activating these neurons and driving increased protein consumption. Additionally, ethanolamine and phosphoethanolamine serve as nitrogen donors. With the assistance of the microbiome, nitrogen from ethanolamine is extracted and incorporated into the biosynthesis of both essential and non-essential amino acids. In fly cells, the ethanolamine-phosphate phospho-lyase (ETNPPL)-dependent pathway facilitates nitrogen extraction from phosphoethanolamine, contributing primarily to the synthesis of non-essential amino acids. Together, these pathways help maintain protein homeostasis under protein-limited conditions. Two-way ANOVA with the Geisser-Greehouse correction, followed by Šidák’s multiple comparisons test, was used to analyze the data in Figures 5A**-5F**. See also **Figure S5**.

These results indicate that these pathways play a critical role in supplying flies with amino acids needed for egg laying, both on the complete and the amino acid-free holidic diet. In germ-free flies, additional factors beyond protein likely also contribute to the reduced egg laying. Together, these findings suggest that nitrogen utilization from the phospholipid reservoir is crucial for supporting reproduction, when dietary amino acids are absent or not provided in adequate amounts.

Collectively, as summarized in the model in **Figure 5G**, our studies describe a previously undocumented nitrogen flow from phospholipids to amino acids, which is enhanced in the absence of sufficient dietary protein. We elucidate the pathways mediating this nitrogen flow and investigate its role in maintaining protein homeostasis. Specifically, protein starvation increases the degradation of phospholipid PE, through phospholipases like PLD, releasing ethanolamine and phosphoethanolamine. Ethanolamine signals protein scarcity to the brain by activating DA-WED protein hunger neurons, leading to increased protein consumption. To restore protein balance, ethanolamine and phosphoethanolamine also act as nitrogen carriers circulating throughout the body. Facilitated by the gut microbiome and the eukaryotic catabolic enzyme ETNPPL, nitrogen is extracted from these molecules and incorporated into amino acid and protein biosynthesis. Thus, phospholipids serve as a hidden nitrogen reservoir, and failure to mobilize nitrogen from this reservoir significantly impacts reproductivity. Together, our findings uncover a previously uncharacterized metabolic plasticity that enables nitrogen redistribution to maintain protein homeostasis.

## Discussion

Protein stands out among macronutrients due to its unique nitrogen content and the body’s inability to biosynthesize essential amino acids. Unlike fats and carbohydrates, the body is generally thought to lack dedicated storage sites for excess protein. However, our research in *Drosophila* reveals that the phospholipid phosphatidylethanolamine (PE) acts as a hidden nitrogen reservoir, storing nitrogen within its headgroup, ethanolamine. When dietary protein is insufficient, nitrogen from this reservoir is mobilized by releasing ethanolamine and its phosphorylated form, phosphoethanolamine, which are then utilized for amino acid and protein synthesis. This process is facilitated by the microbiome and the ETNPPL-dependent pathway in fly cells, which convert ethanolamine and phosphoethanolamine preferentially into essential and non-essential amino acids, respectively. Intriguingly, ethanolamine serves a dual function: in addition to its metabolic role, ethanolamine produced in brain glial cells by PLD acts as a neuromodulator, activating protein-specific hunger neurons that drive the organism to seek and consume protein-rich food. The other metabolic product of PLD-mediated PE degradation, phosphatidic acid (PA), is also a well-established signaling molecule involved in various signaling processes (Zhou et al 2024). Whether PA exerts additional roles by modulating the activity of other neural circuits remains to be determined.

### Temporal dynamics of nitrogen mobilization and protein appetite

The upregulated PLD expression in peripheral tissues, which presumably leads to PE degradation and increased levels of ethanolamine and phosphoethanolamine, peaks as early as two days after protein deprivation (**Figures 3H-3K**). Consistently, circulating phosphoethanolamine levels peak on day 2 (**Figures 1G and 1H**), and ethanolamine levels in virgin females follow a similar pattern (**Figure 1I**). In contrast to this early mobilization of nitrogen, the behavioral switch toward protein preference does not peak until days 5-7 (**Figures 1B-1D**), revealing a temporal delay between metabolic nitrogen release and the onset of protein-seeking behavior. This suggests that flies mobilize nitrogen from the phospholipid reservoir well before they are driven to actively consume protein-rich foods. This early nitrogen mobilization may act as a buffer, supplying nitrogen for amino acid biosynthesis and delaying the immediate need for dietary protein. Such a buffering system could provide flies with additional time to locate and access protein-rich resources, ensuring survival under fluctuating nutrient availability. The delayed onset of protein-specific hunger may also help explain why protein preference can sometimes appear weak or inconsistent in flies and other model organisms, and is therefore easily overlooked.

The mechanisms underlying this delay, however, remain unknown. One possibility is that the brain requires ethanolamine to accumulate beyond a certain threshold to activate DA-WED neurons. Alternatively, the delayed brain response may be regulated by other proteins or cellular processes. Given that CTL1 and CTL2 serve as key ethanolamine transporters mediating its neuromodulatory effects on DA-WED neurons, they may act as gatekeepers controlling the activation of protein appetite. Future studies should explore the specific downstream pathways triggered by CTL1/CTL2 in DA-WED neurons and how they drive protein-seeking behavior.

Understanding these mechanisms will be crucial for deciphering the complex interplay between metabolic nitrogen sensing and behavioral adaptation.

### Baseline nitrogen flow supports processes with high protein demand

While we identified phospholipids as a nitrogen reservoir in the context of protein starvation, our findings suggest the existence of a baseline level of nitrogen flow from phospholipids to amino acids. This is evidenced by our isotope tracing experiments (**Figure 4B**), which show nitrogen incorporation from ethanolamine into amino acids, even in the presence of dietary protein. This implies that nitrogen utilization from phospholipid reservoirs occurs in tissues or life stages with high protein demand, contributing to normal physiological functions. Supporting this, egg-laying—a process heavily dependent on protein—was significantly reduced when nitrogen utilization from the PE reservoir was impaired, even on a complete holidic diet (**Figures 5A–5C**).

Despite our demonstration of the nitrogen flow, the precise cellular and tissue-specific sources of PE remain unknown. A key question is which cellular compartment, membrane type, or tissue reservoir donates PE for degradation without compromising essential cellular functions. Given that PE is a major component of various membranes, its mobilization must be tightly regulated to prevent disruptions in membrane integrity and organelle function. Is there a dedicated subcellular structure responsible for PE degradation and ethanolamine production?

Alternatively, do specific subpopulations of organelles, such as the Golgi, endoplasmic reticulum, lysosome, or endosomes, selectively sacrifice portions of their membranes for PE breakdown under protein starvation conditions? Identifying the precise subcellular and tissue-specific sources of PE and the regulatory mechanisms controlling its degradation will be critical for understanding how cells mobilize membrane lipids to buffer nitrogen scarcity while maintaining structural and functional homeostasis.

### Evolutionarily conserved pathways in nitrogen metabolism

The microbiome is previously described to provide the host with essential amino acids, supporting nutrient acquisition and metabolic health in both insects and humans (Douglas & Prosser 1992, Henriques et al 2020, Kim et al 2021, Leitao-Goncalves et al 2017, Metges 2000). However, the specific nitrogen sources that the microbiome utilizes to synthesize and supply the host with amino acids remain unclear. Here, we provide evidence that ethanolamine serves as one such source. The molecular pathways we identified here, including CTL1/2, PLD, and ETNPPL, are evolutionarily well-conserved. This conservation suggests the possibility of a similar nitrogen recycling mechanism in mammals, allowing animals to utilize nitrogen from phospholipids when dietary protein intake is insufficient, thereby promoting metabolic resilience. In support of this, studies in rats have suggested that ETNPPL expression can be modulated by dietary protein content but not fat (Fleshood & Pitot 1970b). Newborn rats and human infants exhibit significantly higher serum ethanolamine levels compared to adults, correlating with their increased protein demands for growth and development (Dickinson et al 1965, Kume et al 2006). This increase in ethanolamine during anabolic states aligns with its potential role as a nitrogen source during periods of high protein needs. Interestingly, cancer cells have also been shown to elevate phosphoethanolamine levels in response to starvation of the amino acid glutamine, which stimulates tolerance to nutrient starvation and tumor growth (Osawa et al 2019). This metabolic reprogramming highlights a potential parallel mechanism by which cancer cells, like developing tissues, mobilize alternative nitrogen sources to maintain biosynthetic and metabolic activities under nutrient stress. Future studies will be essential to further elucidate the extent to which these nitrogen recycling pathways contribute to metabolic resilience across different physiological and pathological contexts.

### Conclusions

Our study uncovers a previously undocumented metabolic plasticity that animals leverage to cope with dietary protein insufficiency. The molecular pathways identified here provide an opportunity to test the generalization of these mechanisms in other model systems. Several important questions remain for future investigations. For instance, what are the upstream signals that initiate the nitrogen flow by promoting PE degradation? Where is the to-be-degraded PE located—does a dedicated, readily degradable PE pool exist within cells? How is the mobilized nitrogen subsequently utilized? Additionally, is the release of ethanolamine a cell-autonomous process, with each cell orchestrating its own nitrogen mobilization, or do specific cell types play a predominant role in releasing ethanolamine, which is then taken up and utilized by other cells?

Collectively, our findings highlight the remarkable metabolic flexibility animals employ to meet protein demands under nutrient-limited conditions, paving the way for future studies to unravel the regulation and coordination of this previously unrecognized nitrogen reservoir.

## ACKNOWLEDGMENTS

We thank Yuh-Nung Jan, Mark Wu, P. Raghu, Bloomington *Drosophila* Stock Center, and Vienna *Drosophila* Resource Center for the reagents. We thank John Schulze in the Proteomics Core of the Genome Center at the University of California, Davis for the amino acid analysis. We thank Kit-sum Leung from the Stable Isotope Laboratory at the University of Hong Kong and Chris Yarnes from UC Davis Stable Isotope Facility for compound-specific stable isotope analysis using GC-MS. We thank Duncan Holbrook-Smith at the General Metabolics for compound-specific stable isotope analysis using LC-MS. We thank Jesse Nippet from the Stable isotope mass spectrometry lab at Kansas State University for EA-IRMS analysis. We thank Kaveh Ashrafi, Sha Liu, Rushika Perera, Masashi Tabuchi, Peter Turnbaugh, John Vaughen, and members of the Liu lab for helpful discussions. Schematics were created with BioRender.com.

Q.L. acknowledges support from the Edward Mallinckrodt Jr. Foundation, the Esther A. & Joseph Klingenstein Foundation, the Pew Scholars Program in the Biomedical Sciences, and the Program for Breakthrough Biological Research. This work was supported by NIH grant NIGMS R35GM138060 (Q.L.).

## AUTHOR CONTRIBUTIONS

Q.L. and T.M. conceived the project. T.M. carried out behavioral analyses and prepared samples for amino acid and isotope analyses. Q.L. and C.S. established the plate assay. G.W. performed electrophysiology experiments. T.M., G.W., and L.T. performed immunostaining. M.P-W., S.V., N.A., C.S., A.T., and R.W.S helped with the feeding behavioral analysis. T.M. and P.M.E. performed the sample preparation for LC-MS. T.M. and Q.L. analyzed the data and wrote the manuscript with input from all authors.

## Materials and Methods

### Fly Stocks

The wild-type *iso^31^* and *TH-F3-Gal4* were obtained from M. Wu (Johns Hopkins University, MD). *pld3.1* mutant line was provided by P. Raghu (National Centre for Biological Sciences, India), *w^1118^* wild-type was provided by Y. Jan (University of California, San Francisco, San Francisco), and *EAS^KO^* was provided by J. Steinhauer (Yeshiva University, NY). *UAS-CTL1 RNAi* (VDRC KK102538), *UAS-CTL2 RNAi* (VDRC KK100290), PLD-Tagged line (VDRC #318697), PLD TRiP RNAi (BDSC #32839), *PLD RNAi* (VDRC KK102796), *Tub-Gal4* (VDRC #67048)*, NP1-Gal4* (generated from BDSC #67057), *ETNPPL* background control line MI04832 (BDSC #38587), ETNPPL mutant line MI15731 (BDSC #61139), were obtained from the Bloomington *Drosophila* Stock Center (BDSC) or the Vienna *Drosophila* Resource Center (VDRC). All transgenic lines used in this study were back-crossed at least five times to *iso^31^* wild-type prior to experiments, with the exception of *ETNPPL* background control line MI04832, ETNPPL mutant line MI15731, *w^1118^* wild-type, and *EAS^KO^*.

### Fly husbandry

Flies were maintained on a standard molasses diet (889mL water, 10g agar, 15g yeast, 60g cornmeal, 80mL molasses, 8mL tegosept, 2.4mL propionic acid, and 0.28mL phosphoric acid) in 12-hour light:12-hour dark cycles at 25°C and 60% humidity unless indicated otherwise.

Chemically defined holidic media were prepared according to previous protocols (Piper et al 2014). Briefly, agar, sucrose, branched-chain amino acids, and amino acids with low solubility (L-Leucine, L-Isoleucine, L-Tyrosine) were added to solutions containing metal ions and cholesterol. The mixtures were autoclaved for 15 minutes at 121°C. Filter-sterilized acetate buffer and solutions of the remaining amino acids, lipids, vitamins, nucleotides, inositol, choline, and preservatives were added while mixtures were stirred on a hot plate not to exceed 65°C. After mixing, 3-5mL of food was dispensed into vials and stored at 4°C until use, but no longer than 3 weeks.

For the two-choice plate feeding assay, immunostaining, amino acids analysis, isotope tracing experiments, and egg-laying assay, 0-2 days old male and virgin female flies of the appropriate genotype were collected under CO2 anesthesia and allowed to recover for 2-3 days, before being transferred to holidic food with or without amino acids, or with different supplementations. Age-matched mated female flies were obtained by co-housing virgin females with wild-type *iso31* males during the recovery period.

### Single fly two-choice plate feeding assay

Flies were collected under CO_2_ anesthesia within 3 days after eclosion and allowed to recover on either a standard molasses diet or a holidic diet for at least 2 days in 12-hour light:12-hour dark cycles at 25°C and 60% humidity before the assay. Two 96-well cell culture plates were filled with 7.5% agar mixed with either 200 mM sucrose and 2% blue dye (FD&C #1) or 10% yeast and 6% red dye (FD&C #40). Flies were cold-anesthetized on ice for 10 minutes and individually loaded to an ice-cold empty 96-well plate before being flipped into one food plate. The second food plate was then aligned and secured with a rubber band, forming an apparatus with 96 independent feeding chambers. Since the loading step takes longer than flipping the flies onto the food plate, using an ice-cold empty plate helps minimize fly activity during loading, ensures equal exposure time for both plates, and maintains their initial temperature at room temperature. To minimize gravitational bias, the apparatus was positioned vertically, ensuring that both food sources were at the same horizontal level. The setup was then transferred to a black box to block the light and placed in a 25°C incubator for 2 hours of feeding. After feeding, flies were sacrificed by freezing at -20°C for at least 30 minutes. They were then transferred to an empty plate and heated for 10 minutes on an 80°C hot plate before being crushed using a 3D-printed 96-well fitted smasher. Large fly body fragments were manually removed. The extracted dye was quantified at 628 nm (blue dye for sucrose) and 505 nm (red dye for yeast) using a plate reader. Switching the color association with yeast and sucrose did not affect the results. The amount of food consumed from each source was calculated using standard curves generated from serial dye dilutions. Protein preference was calculated using the following formula:

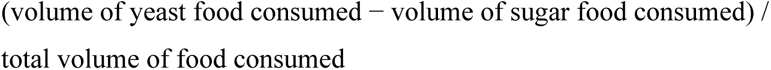

### Hemolymph collection and amino acid analysis

For hemolymph collection, around 200 flies were used for each condition per replicate. Single-edge blades were used to separate the head and body on a pre-iced agar pad. Immediately after decapitation, the body was gently squeezed with forceps, releasing a droplet of hemolymph, which was collected via 0.5μl micropipettes and transferred to a PCR tube on ice. A total of 3-4 μl was collected per condition and sent for amino acid analysis at the UC Davis Molecular Structure Facility. The facility utilizes a Hitachi LA8080 amino acid analyzer, which operates with a lithium citrate buffer system optimized for physiological hemolymph samples. Ion-exchange chromatography was used to separate amino acids, followed by post-column ninhydrin reaction detection, for the quantification of individual amino acids.

### Dissection and Immunostaining

Brains were dissected by removing the head cuticle and clearing the surrounding trachea. For fat body dissection, the dorsal cuticle with attached abdominal fat body was separated from the ventral cuticle and underlying tissues using forceps. The gut was dissected by removing the head and opening the abdomen, allowing the entire gut tissue to be carefully extracted. Dissected tissues were fixed in 4% paraformaldehyde (PFA) for 30–40 minutes at room temperature, then washed in PBST (PBS + 0.3% Triton X-100) and blocked with 5% normal goat serum (NGS) in PBST for 1 hour at room temperature. Samples were incubated in darkness with primary antibodies (mouse anti-V5, Invitrogen, 1:500) at 4°C for 16–40 hours, followed by washing and incubation with secondary antibodies Alexa 488 anti-mouse (Invitrogen, 1:1000) or Alexa 568 anti-mouse (Invitrogen, 1:1000). DAPI (Invitrogen No. D3571, 1:1000) was diluted in PBS and incubated for 10–15 minutes at room temperature for nuclear staining. Samples were then transferred to PBS with 70% glycerol for transparency and mounted using mounting medium (Fisher, H-1000NB). For native GFP fluorescence, brains were fixed in 4% PFA for 5 minutes, washed three times in PBST (10 minutes each), and then incubated in PBS with 70% glycerol and mounted. All procedures with native GFP were performed in darkness, with samples protected from light using aluminum foil coverings. All Imaging was obtained using a Zeiss LSM700 confocal microscope, capturing 1 µm thick sections at 25x or 63x magnification.

### PLD-GFP-V5 Quantification

To quantify V5-tagged PLD expression in the gut and fat body in **Figures 3H-3K** and **S3B-S3C**, the number of puncta and total puncta volume were analyzed. Images were taken under 63x magnification and analyzed in Fiji. Image stacks containing PLD-V5 expression were extracted from the original stacks. The puncta number and total puncta volume were automatically calculated using the “*3D Objects Counter”* plugin, with a consistent threshold value applied to all samples.

### Generation of germ-free flies

The protocol to generate axenic *iso^31^* fly cultures by sterilizing embryos was adapted from Leitao-Goncalves et al. (2017) (Leitao-Goncalves et al 2017). Adult *iso^31^* flies (∼5 days old) were collected in an inverted 250 mL beaker, sealed with a cell culture petri dish containing 10 g yeast, 10 g glucose, 1 g agar, 80 mL H₂O, and 20 mL apple juice, with the surface rinsed in a yeast solution. After incubating for 4–5 hours at 25°C, embryos were collected from the petri dish and sterilized by sequential treatments: 2.5% active chlorine (50% bleach) for 2 min, followed by 70% ethanol for 2 min, and autoclaved distilled water for 2 min. The embryos were then transferred onto sterile food (autoclaved before being poured into culture vials) supplemented with antibiotics at the following final concentrations: 416.7 μg/mL tetracycline (high dose), 41.67 μg/mL chloramphenicol, 41.67 μg/mL ampicillin, and 8.333 μg/mL erythromycin. The yeast content of the medium was increased to 41.67 g/L to compensate for the developmental delay.

The absence of bacteria was assessed by grinding flies in sterile 1× PBS and spreading the suspension on LB and MRS plates, which were incubated at 37°C before checking for bacterial colony formation. Additionally, bacterial contamination was evaluated using 16S rDNA PCR.

### Stable isotope analysis of ^15^N labeled amino acids

#### Gas chromatography-isotope ratio mass spectrometry (GC-IRMS)

^15^N-amino acids from ^15^N-ethanolamine-fed animals were analyzed by compound specific stable isotope analysis, using gas chromatography-isotope ratio mass spectrometry (GC-IRMS) (**Figures 4B-4C, 4G, 4I, and S4A-S4F**). Flies were fed on a ^15^N-ethanolamine-supplemented diet for 5 days, being flipped into fresh food every 2-3 days. After the feeding period, flies were transferred into glass vials and dried in a 65°C incubator for at least 4 hours. Dried samples were then cooled to room temperature before being shipped for analysis. GC-IRMS analyses were performed at two facilities: the Stable Isotope Facility, Department of Plant Sciences, University of California, Davis, and the Stable Isotope Mass Spectrometry Laboratory, University of Hong Kong. Both facilities conducted GC-IRMS using the same model of instrument and yielded similar results. The protocols from both facilities are similar and are briefly described below.

For the UC Davis facility, amino acids were liberated from proteins by acid hydrolysis (6 M HCl, 70 min, 150°C under an N₂ headspace). Amino acids were then derivatized as *N*-acetyl methyl esters (NACME) to render them suitable for gas chromatography (GC). Additional purification steps, such as strong cation-exchange chromatography (SCX) using Dowex 50WX8 resin, may be required for samples with significant fractions of carbohydrates, lipids, salts, or other potential matrix interferences. NACME amino acid derivatives were injected at 260°C (splitless, 1 min) and separated on an Agilent DB-35 column (60 m × 0.32 mm ID × 1.5 μm film thickness) at a constant flow rate of 2 mL/min under the following temperature program: 70°C (hold 2 min); 140°C (15°C/min, hold 4 min); 240°C (12°C/min, hold 5 min); and 255°C (8°C/min, hold 35 min). GC-IRMS was performed on a Thermo Trace GC 1310 gas chromatograph coupled to a Thermo Scientific Delta V Advantage isotope-ratio mass spectrometer via a GC IsoLink II combustion interface. The combustion reactor consisted of a NiO tube containing CuO and NiO wires maintained at 1000°C. Water was subsequently removed through a Nafion dryer before the analyte gases were transferred to the IRMS. During the ^15^N analysis, CO₂ was removed from the post-combustion carrier stream using a liquid nitrogen trap to prevent isobaric interferences within the ion source.

For the University of Hong Kong facility, isotope analysis was also performed using a Thermo GC Trace 1310 gas chromatograph connected to a Thermo IRMS Delta V Advantage. Samples were first lipid-extracted by adding 10 mL of Folch solution (chloroform:methanol, 2:1 v/v) to the sample. The samples were homogenized by vortexing, sealed with nitrogen gas, and incubated horizontally in an ice box on a shaker for 30 min. The supernatant containing the extracted lipids was removed, and the fat-extracted sample was dried at room temperature in a hood. Acid hydrolysis was then performed by adding 1 mL of ultra-clean 6 M HCl and incubating at 110°C for 20 hours. Carboxyl derivatization was carried out using a mixture of acetyl chloride and isopropanol (1:4) with heating at 110°C for 1 hour. Amine derivatization was performed by adding 0.5 mL of dichloromethane (DCM) and 0.5 mL of trifluoroacetic anhydride (TFA anhydride) to each sample vial, heating at 110°C for 10 minutes, and concentrating the sample using a nitrogen evaporator before instrumental analysis.

#### Liquid chromatography-mass spectrometry (LC-MS)

To distinguish the different heavy isoforms of amino acids (**Figures 4D–4E**), liquid chromatography-mass spectrometry (LC-MS) was performed by General Metabolics, LLC, Boston. Whole-body fly samples were first acid-hydrolyzed with 6N HCl at 110°C for 20 hours to liberate amino acids from proteins. The samples were then frozen at -80°C for at least 24 hours and subsequently freeze-dried using a vacuum pump for at least 24 hours until completely dry before being shipped for analysis. Samples were analyzed in positive ionization mode, with analytes separated using hydrophilic interaction liquid chromatography (HILIC). MS1 mass spectra were collected using high-resolution mass spectrometry. Quality control (QC) samples, composed of a standard metabolite mix, were measured to confirm system suitability and retention time stability. LC-MS detected most of the canonical amino acids and their labeled forms. The intensity for each isotopologue of the measured metabolites is presented both as raw ion intensity and as a mass distribution vector (MDV), which describes the fraction of the detected metabolite that corresponds to each labeled form.

#### Elemental analyzer-isotope ratio mass spectrometry (EA-IRMS)

For *iso^31^* wild-type head and body samples (**Figure S4H**), bulk analysis of ^15^N elemental composition was performed at the Stable Isotope Mass Spectrometry Laboratory at Kansas State University, using an Elementar vario Pyro cube coupled to an Elementar Vision mass spectrometer. Samples were first dried and homogenized into a fine powder before being shipped for analysis. During analysis, samples were combusted at high temperatures to convert nitrogen-containing compounds into gases for isotope ratio measurement.

### Egg-laying Assay

Following eclosion, 5–6 virgin females were collected into a standard molasses food vial with 2 males to allow mating for 48 hours. Males were then removed, and mated females were transferred to either a complete or an amino acid-free holidic diet. Mated females were flipped into a fresh food vial every 24 hours, and eggs from the previous vial were counted. For rescue experiments, yeast was added during the 48-hour mating period and also supplemented in the vials during the egg-laying process.

### Perforated patch-clamp recordings

Flies were anesthetized on ice, and then brains were dissected in the pre-bubbled with 95% O_2_ and 5% CO_2_ saline solution, containing 101 mM NaCl, 3 mM KCl, 1 mM CaCl_2_, 4 mM MgCl_2_, 1.25 mM NaH_2_PO4, 20.7 mM NaHCO_3_, 5 mM glucose, and 10 mM HEPES (pH adjusted to 7.2). To better visualize the cell body of DA-WED neurons, the perineuronal sheath surrounding the brain was first focally and carefully removed, and then a mixture of collagenase (0.2 mg/mL) and dispase (0.8 mg/mL) was used to treat brains at room temperature for about 3-5 min with variable time length based on the consequent performance of DA-WED cell bodies. According to the accessibility of the recording site observed under the microscope and to improve the subsequent successful ratio as much as possible, additional cleaning steps may be applied through using a small stream of saline pressure ejected from a pipette with a 0.5 mL syringe connected to the pipette holder. Brains were discarded if electrodes could not access the cell body ideally under microscope. Ideal brain preparations with posterior facing up were then placed at the bottom center of the circular recording chamber with one glass coverslip sealed as the bottom and immobilized using a custom-made platinum anchor in a V shape. The recording chamber was placed on an X-Y stage platform of motorized movable base plate (Scientifica), and the cell bodies of the target neurons were visualized with GFP fluorescence on a fixed-stage upright slice scope and viewed with camera capture of a 40x water immersion objective lens. Visualization was achieved by IR-DIC optics when the recording electrode approached the cell bodies of target neurons.

Recording electrode pipettes were made from borosilicate glass capillary (without filament) by the P1000 puller (Sutter Instruments), and further polished with a microforge (MF-200, WPI) before recording to improve the successful ratio. Internal pipette solutions were made from 102 mM potassium gluconate, 0.085 mM CaCl_2_, 0.94 mM EGTA, 8.5 mM HEPES, 4 mM Mg-ATP, 0.5mM Na-GTP, 17 mM NaCl with pH adjusted to 7.2, which was then filter sterilized and aliquoted into 1 mL tube to be stored in -20°C. The chemical compound β-escin was used to form the perforated whole-cell configuration, which was prepared and aliquoted as a 50 mM stock solution in water and stored up to 2 weeks at -20°C. Before experiments, β-escin was taken out and thawed at room temperature, and then was added into the internal pipette solution to get a final concentration of 50 μM. As β-escin is light-sensitive, the internal pipette solutions after the addition of β-escin were covered with aluminum foil. When filling recording pipettes, pipette tips were first dipped quickly for 1 sec in the internal pipette solution before being back-filled using the microloader tips (Eppendorf). Usually there are many air bubbles inside the pipettes, and they must be removed first by gentle tapping. β-escin in the internal solutions remained efficient and stable for several hours without evidence of the formation of precipitations. For application of ethanolamine, it was always freshly added into the internal recording solutions to obtain a final concentration of 0.5 mM. For application of phosphoethanolamine, it was first made to a concentration of 100 mM, and then was added to the internal solutions to obtain indicated final target concentrations. All these compounds were perfused to the recording chamber to act on DA-WED neurons through the peristaltic perfusion system (Golander pump).

The multiclamp 700B amplifier (Molecular Devices) connected to the Axon Digidata 1550B (Molecular Devices) was used to perform all electrophysiological experiments controlled by the computer system (Dell). pCLAMP 11.1 software package was used for operating these devices and for data acquisition. The acquired signals were sampled at 20 kHz and low-pass filtered at 2 kHz. Junction potentials were corrected prior to the formation of high-resistance (GΩ) seal. After filling the internal pipette solution, the pipette had a variable 9-12 MΩ resistance with negative pressure given through one custom-made tubing. After confirming the recording site with adjusted focus, the recording electrode was slowly moved into the saline solution and toward the target neuron through hand using the micromanipulator (Scientifica). After observing clear contact between the pipettes and the cell body, release of the negative pressure inside the pipettes generally could help get a high resistance sealing up to GΩ, applying the negative pressure to facilitate the sealing if necessary. If this sealing failed with low sealing resistance, the neuron would be disregarded, and a second attempt would be made on another corresponding neuron in the other brain hemisphere. If the second attempt failed again, the brain would be discarded. After a successful GΩ sealing was established, the perforated whole-cell patch clamp model was allowed to develop spontaneously and slowly over time (8-15 minutes) without any negative pressure suction applied inside the pipette. Experiments were terminated if perforated patches were not formed after 15 minutes. As the breakthrough became evident, indicated by the gradual development of a large capacitance transient in the seal test window of pCLAMP 11.1, access resistance was monitored using the membrane test function from the formation of high resistance to the completion of the perforation process. If the access resistance was less than 40 MΩ when neurons reached a steady state, data recording would be executed. All electrophysiological recordings were conducted at room temperature.

### Data plotting and Statistics

Data plotting and Statistical analyses were performed using Prism software (GraphPad). For normally distributed datasets, scattered dot plots with mean ± SEM are shown. For non-normally distributed datasets, box plots are shown with scattered dots to display all raw data points. The line inside the box indicates the median. The top and bottom of the box represent the 75th and 25th percentiles, respectively. The whiskers represent the minimum and maximum values. To visualize percentage changes, heat maps were used, where blue indicates a reduction, red indicates an increase, and white indicates no change. For electrophysiological changes before and after ethanolamine supplementation, connected dot plots were used.

For comparisons of 2 groups of normally distributed data, unpaired or paired t-tests with or without Welch’s correction were performed for data sets with or without differences in variances, respectively. For comparisons of 2 groups of non-normally distributed data Mann-Whitney tests were performed. For multiple comparisons of normally distributed data with no differences in variance, ANOVAs followed by Tukey multiple comparisons were performed. For multiple comparisons of normally distributed data with different variances, Brown-Forsythe and Welch ANOVA tests were performed followed by post hoc multiple comparisons. For multiple comparisons of non-normally distributed data, Kruskal-Wallis tests were performed, with Bonferroni correction for Dunn’s multiple comparisons test.

---

**Figure S1.**
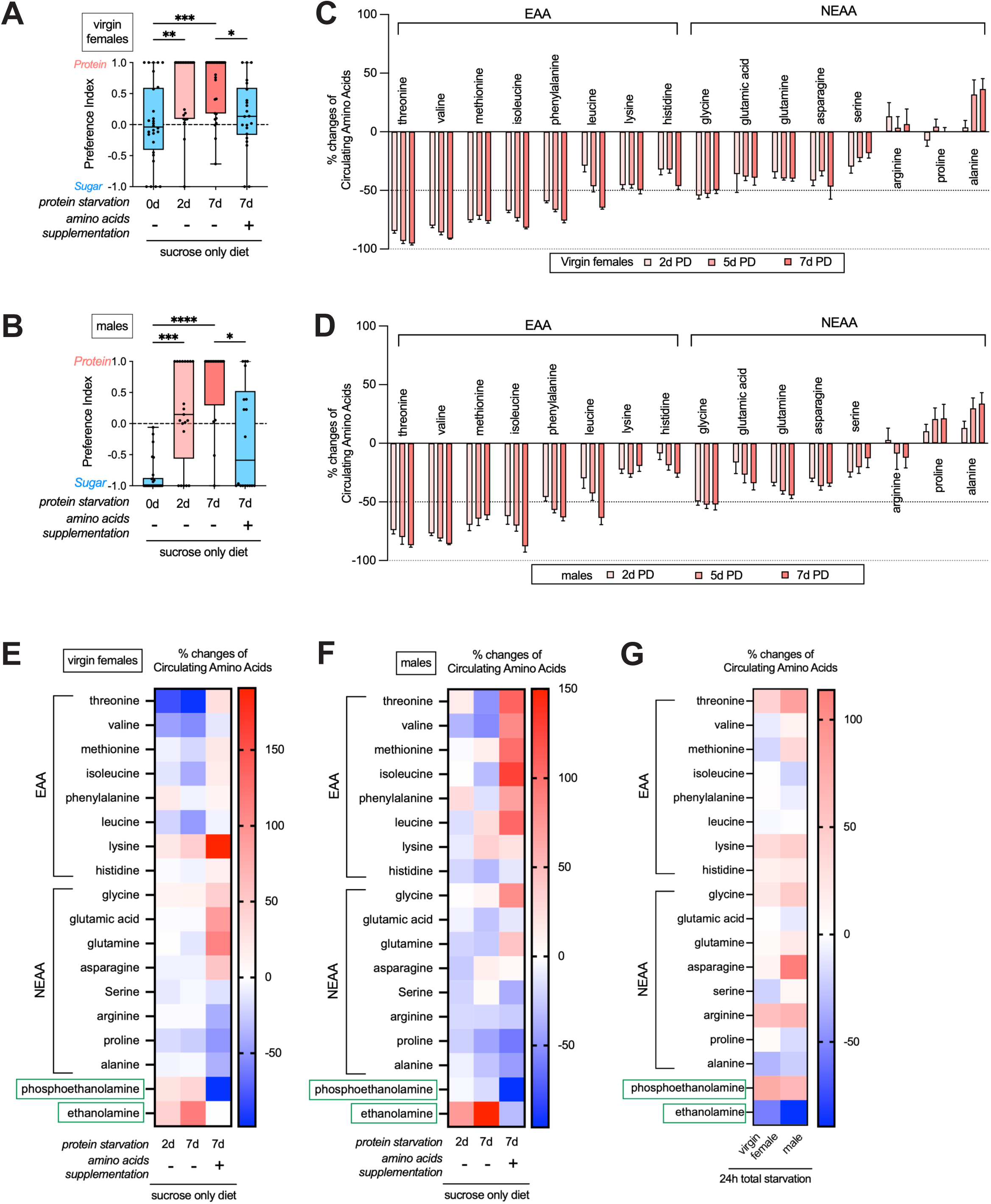
Protein starvation raises circulating ethanolamine and phosphoethanolamine, related to Figure 1. (**A-B**) Protein preference index in age-matched wild-type *iso31* virgin female (**A**) and male (**B**) flies subjected to protein starvation on a sucrose-only diet for 0, 2, and 7 days with or without amino acids supplementation. n=21-29 in **A**, n=16-26 in **B**. (**C-D**) The same dataset as in Figures 1E an**d 1F**, showing bar graph plot of the % changes in circulating amino acids from the hemolymph of age-matched wild-type virgin female (**C**) and male (**D**) flies following protein deprivation (PD) on an amino acid-free holidic diet for 2, 5, and 7 days, compared to control flies without protein starvation. n=7 replicates in **C** and 8 in **D**. Each replicate contains hemolymph collected from at least 200 flies per condition. (**E-F**) % changes in circulating amino acids from the hemolymph of age-matched wild-type virgin female (**E**) and male (**F**) flies following protein starvation on a sucrose-only diet for 0, 2, and 7 days with or without amino acids supplementation. n=7 replicates in **E**, n=2-5 replicates in **F.** Each replicate contains hemolymph collected from at least 200 flies per condition. (**G**) % changes in circulating amino acids from the hemolymph of age-matched wild-type virgin female and male flies following complete food starvation for 24 hours on wet kimwipe. n=3 replicates. Each replicate contains hemolymph collected from at least 200 flies per condition. Kruskal-Wallis tests were performed with Dunn’s multiple comparisons test for **Figures S1A** and **S1B**. In this and the subsequent figures, for normally distributed datasets, scattered dot plots with mean ± SEM are shown. For non-normally distributed datasets, box plots are shown with scattered dots to display all raw data points. The line inside the box indicates the median. The top and bottom of the box represent the 75th and 25th percentiles, respectively. The whiskers represent the minimum and maximum values. “*”, “**”, “***”, “****”, and “ns” denote P<0.05, P<0.01, P<0.001, P<0.0001, and not significant, respectively.

**Figure S2.**
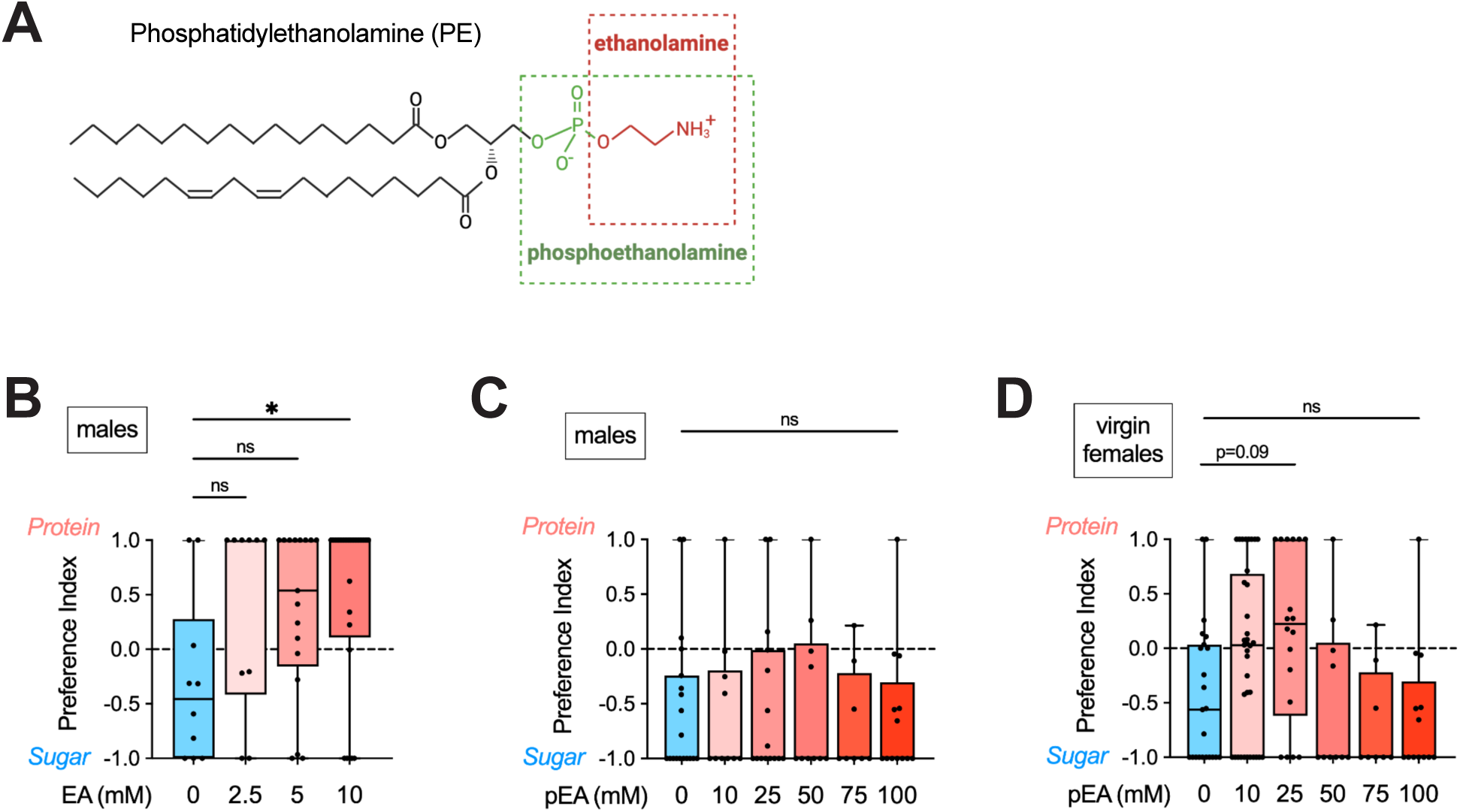
Ethanolamine activates the DA-WED protein hunger neurons, related to Figure 2. (**A**) Structure of phosphatidylethanolamine (PE), with their headgroup, ethanolamine and phosphoethanolamine, highlighted in red and green dash-lined boxes, respectively. (**B**) Protein preference index for age-matched wild-type *iso31* male flies, fed on amino acid-free holidic diets supplemented with 0mM, 2.5mM, 5mM, or 10mM ethanolamine for 3 days prior to two-choice plate feeding assay. Flies were first entrained on a complete holidic diet for 2-3 days before being transferred to ethanolamine-supplemented amino acid-free holidic diets. n=10-25. (**C-D**) Protein preference index for age-matched wild-type *iso31* male (**C**) and virgin female (**D**) flies, fed on amino acid-free (**C**) and complete (**D**) holidic diets supplemented with 0mM, 10mM, 25mM, 50mM, 75mM, or 100mM phosphoethanolamine for 3 days prior to two-choice plate feeding assay. Flies were first entrained on a complete holidic diet for 2-3 days before being transferred to phosphoethanolamine-supplemented diets. n=8-19 for **C** and n=8-36 for **D**. Kruskal-Wallis tests with Dunn’s multiple comparisons test were performed for **Figures S2B-S2D**.

**Figure S3.**
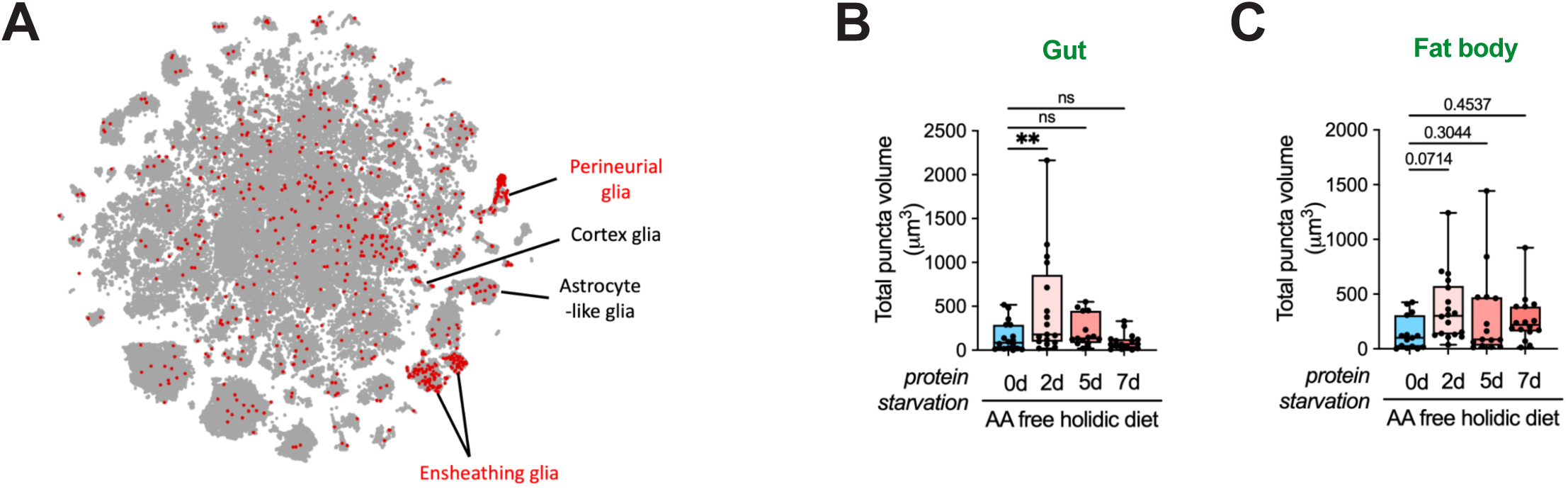
Protein starvation triggers the breakdown of phospholipid PE, related to Figure 3. (**A**) *t-*Distributed stochastic neighbor embedding (*t-*SNE) plot showing the expression of PLD gene in single cells from the wide-type fly brain with major glial cell types annotated. Data were generated using the database from (Dopp et al 2024) (**B-C**) Quantification of total volume of PLD-GFP-V5 puncta immunostained with anti-V5 antibodies in the enterocytes of the R3 gut section (**B**) and fat body cells (**C**) from age-matched virgin female flies subjected to protein starvation on an amino acid-free holidic diet for 0, 2, 5, and 7 days. n=15-18 for **B** and n=22-27 for **C.** Kruskal-Wallis tests with Dunn’s multiple comparisons test were performed for **Figures S3B, S3C.**

**Figure S4.**
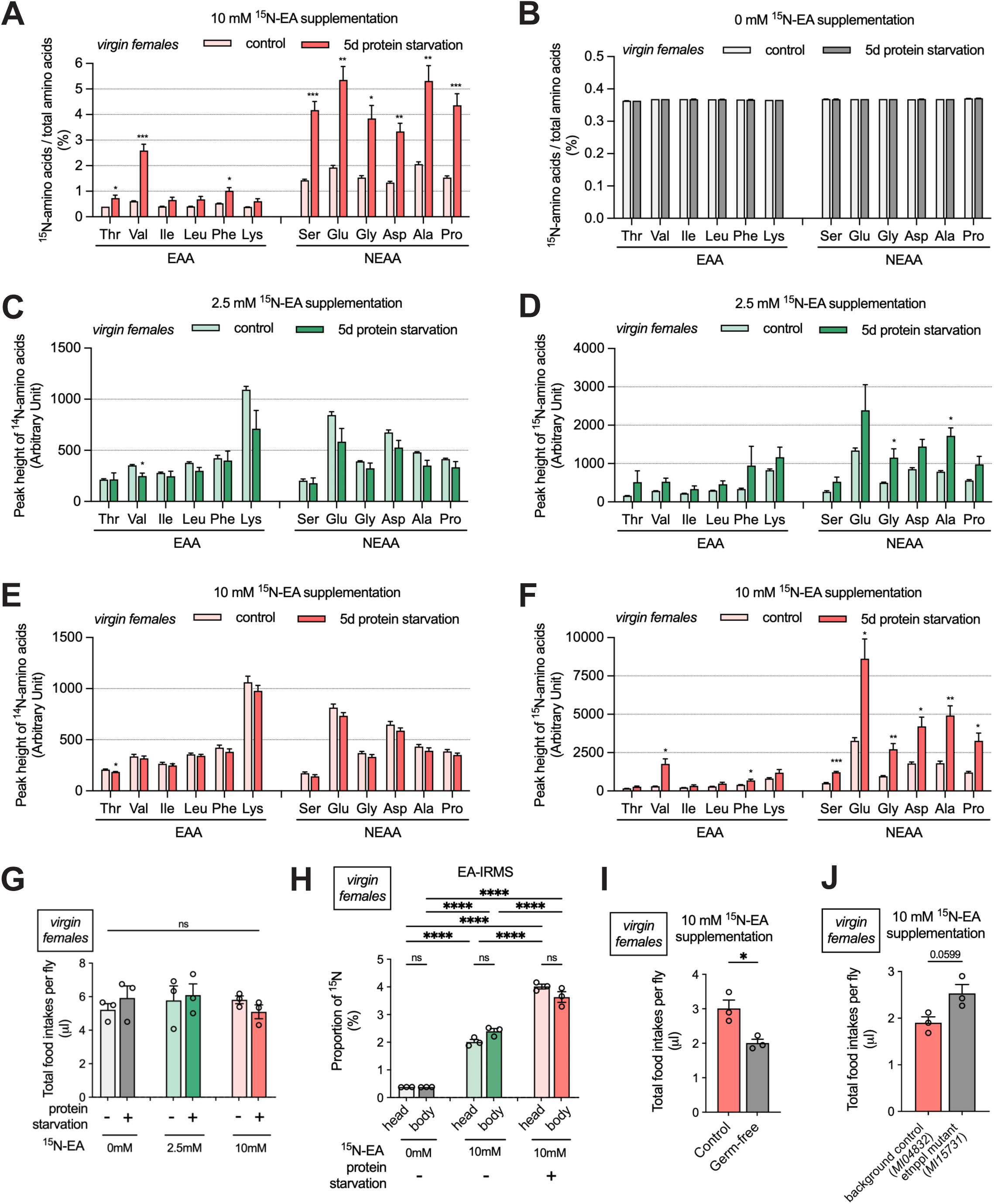
Nitrogen flow from ethanolamine to amino acids, related to Figure 4. (**A**) Percentage of ^15^N-amino acid relative to total amino acids in age-matched wild-type *iso31* virgin female flies fed for 5 days on 10 mM ^15^N-EA supplemented complete holidic diets (control) or amino acid-free holidic diet (5d protein starvation). n= 3 replicates, with each replicate containing 20-30 flies per condition. (**B**) Percentage of ^15^N-amino acid relative to total amino acids in age-matched wild-type *iso31* virgin female flies fed for 5 days on complete holidic diets (control) or amino acid-free holidic diet (5d protein starvation) without ^15^N-EA supplementation. n= 3 replicates, with each replicate containing 20-30 flies per condition. (**C-D**) Peak signal heights of ^14^N-amino acids (**C**) and ^15^N-amino acids (**D**) in age-matched wild-type *iso31* virgin female flies fed for 5 days on 2.5 mM ^15^N-EA supplemented complete holidic diets (control) or amino acid-free holidic diet (5d protein starvation). n= 3 replicates, with each replicate containing 20-30 flies per condition. (**E-F**) Peak signal heights of ^14^N-amino acids (**C**) and ^15^N-amino acids (**D**) in age-matched wild-type *iso31* virgin female flies fed for 5 days on 10 mM ^15^N-EA supplemented complete holidic diets (control) or amino acid-free holidic diet (5d protein starvation). n= 3 replicates, with each replicate containing 20-30 flies per condition. (**G**) Total food intake per fly over 5 days of feeding on ^15^N-EA supplemented holidic diets, with or without amino acids, in age-matched wild-type *iso31* virgin female flies. n= 3 replicates, with each replicate containing 8-16 flies per condition. (**H**) Proportion of ^15^N atoms in head and body samples of wild-type *iso31* virgin female flies fed for 5 days on 0 mM and 10 mM ^15^N-EA supplemented holidic diet with or without amino acids. n= 3 replicates, with each replicate containing 150-200 heads for head samples and 25-30 decapitated bodies for body samples. (**I**) Total food intake per fly over 5 days of feeding on 10 mM ^15^N-EA supplemented complete holidic diets in age-matched control and germ-free virgin female flies. n= 3 replicates, with each replicate containing 8-16 flies per condition. (**J**) Total food intake per fly over 5 days of feeding on 10 mM ^15^N-EA supplemented complete holidic diets in age-matched background control *MI04832* (BDSC_38587) and *etnppl* mutant *MI15731*(BDSC_61139) virgin female flies. n=3 replicates, with each replicate containing 8-16 flies per genotype. Multiple Unpaired t-tests with Welch correction were performed for **Figures S4A-S4F**. ANOVA followed by Tukey’s multiple comparisons test was used to analyze **Figures S4G** and **S4H**. Unpaired t-test was used to analyze data in **Figures S4I** and **S4J.**

**Figure S5.**
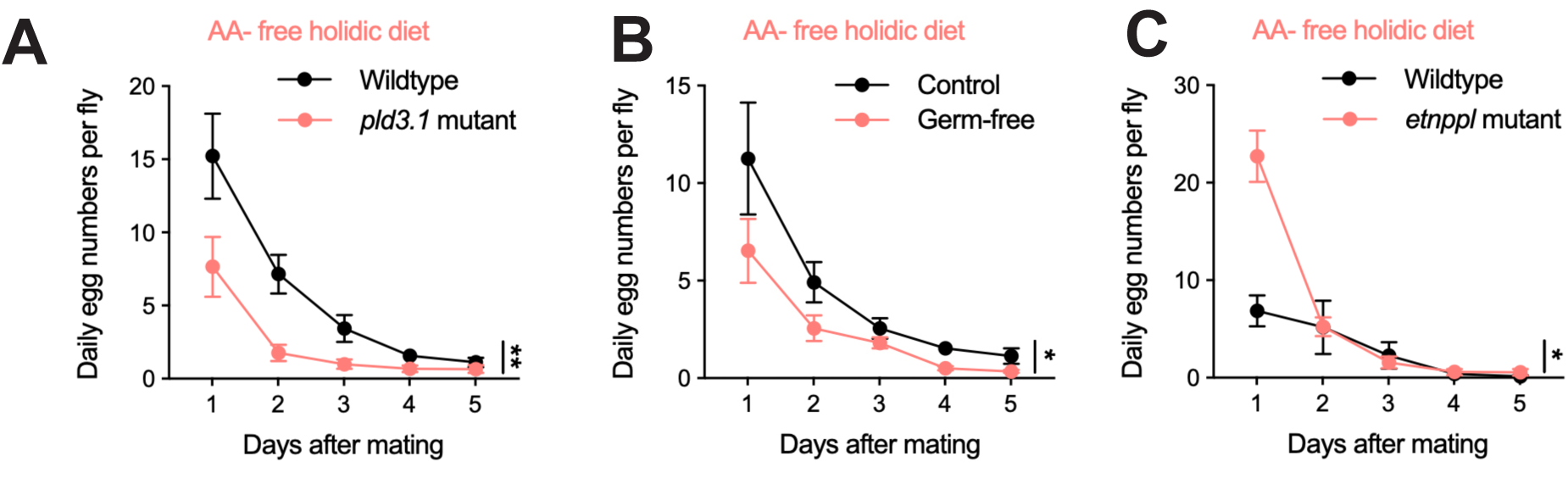
Impaired nitrogen utilization from phospholipids compromises female fertility, related to Figure 5. (**A-C**) Daily egg-laying numbers per fly in age-matched mated female flies of wild-type and *pld 3.1* mutants (**A**), control and germ-free flies (**B**), background control *MI04832* (BDSC_38587) and *etnppl* mutant *MI15731*(BDSC_61139) (**C**) on an amino acid-free holidic diet. n=4-7 replicates, with each replicate consisting of 5-6 virgin female flies mixed with 2 wild-type males. Flies were allowed to mate for 48 hours on a standard molasses diet before females were transferred to an amino acid-free holidic diet. Egg numbers were counted every 24 hours after transferring female flies into a new vial with an amino acid-free holidic diet. Two-way ANOVA with the Geisser-Greehouse correction, followed by Šidák’s multiple comparisons test, was used to analyze the data in **Figures S5A-S5C**.

